# NSAID-mediated cyclooxygenase inhibition disrupts ectodermal derivative formation in axolotl embryos

**DOI:** 10.1101/2024.10.30.621122

**Authors:** Emma J. Marshall, Raneesh Ramarapu, Tess A. Leathers, Nikolas Morrison-Welch, Kathryn Sandberg, Maxim Kawashima, Crystal D. Rogers

## Abstract

Embryonic exposures to non-steroidal anti-inflammatory drugs (NSAIDs) have been linked to preterm birth, neural tube closure defects, abnormal enteric innervation, and craniofacial malformations, potentially due to disrupted neural tube or neural crest (NC) cell development. Naproxen (NPX), a common non-steroidal anti-inflammatory drug (NSAID) used to relieve pain and inflammation, exerts its effects through non-selective cyclooxygenase (COX) inhibition. Our lab has identified that the cyclooxygenase (COX-1 and COX-2) isoenzymes are expressed during the early stages of vertebrate embryonic development, and that global inhibition of COX-1 and COX-2 function disrupts NC cell migration and differentiation in *Ambystoma mexicanum* (axolotl) embryos. NC cells differentiate into various adult tissues including craniofacial cartilage, bone, and neurons in the peripheral and enteric nervous systems. To investigate the specific phenotypic and molecular effects of NPX exposure on NC development and differentiation, and to identify molecular links between COX inhibition and NC derivative anomalies, we exposed late neurula and early tailbud stage axolotl embryos to various concentrations of NPX and performed immunohistochemistry (IHC) for markers of migratory and differentiating NC cells. Our results reveal that NPX exposure impairs the migration of SOX9+ NC cells, leading to abnormal development of craniofacial cartilage structures, including Meckel’s cartilage in the jaw. NPX exposure also alters the expression of markers associated with peripheral and central nervous system (PNS and CNS) development, suggesting concurrent neurodevelopmental changes.

## I. Introduction

Non-steroidal anti-inflammatory drugs (NSAIDs) such as ibuprofen, acetaminophen, and naproxen (NPX) mediate pain and inflammation via inhibition of the cyclooxygenase (COX) isoenzymes. There are two major COX isoforms, COX-1 and COX-2, which produce hormone-like signaling molecules called prostaglandins (PGs) from arachidonic acid isolated from membrane phospholipids (Garavito and DeWitt, 1999). Among other functions, PGs are responsible for generating inflammation and the fever, pain, redness, and swelling that follow inflammatory processes (Ricciotti and FitzGerald, 2011). Despite being well-studied in normal tissue homeostasis (Chen et al., 2019; Goldyne, 2000; Lopez and Ballaz, 2020; Sellers et al., 2010; Vegiopoulos et al., 2010), pathophysiologic inflammation (Coon et al., 2007; Hoozemans et al., 2008; Jackson et al., 2000; Park and Christman, 2006; Sales and Jabbour, 2003; Wang et al., 2005), and tissue healing contexts (Amico et al., 2004; Futagami et al., 2002; Jugdutt, 2007; Radi and Khan, 2005; Takeuchi and Amagase, 2018), much less is known about the role of COX signaling during embryonic development. Additionally, there is contention about the specific roles that COX-1 and COX-2 may play in each of these various contexts and how their roles may determine the specific effects of non-selective NSAIDs versus COX- 2-selective NSAIDs (Pairet and Engelhardt, 1996; Zidar et al., 2009).

In mammalian models, *in utero* exposure to NSAIDs causes congenital defects in craniofacial, cardiovascular, and gastrointestinal tissues (Hultzsch et al., 2021; Nakhai-Pour and Berard, 2008). Work in zebrafish embryos identified that genes encoding COX signaling pathway factors are expressed at the earliest stages of cell specification and that COX signaling is necessary for major developmental processes, including gastrulation (Cha et al., 2005; Cha et al., 2006b). In zebrafish, mouse, and chicken embryo models, COX inhibition via ibuprofen exposure during early development caused abnormal colonization of the gut by enteric nerves derived from neural crest (NC) cells (Schill et al., 2016). Additionally, COX signaling inhibition in chicken embryos altered the expression of factors involved in key pathways for NC cell development and migration including the Wnt/β-catenin and TGF-β pathways (Parmar et al., 2021). Conversely, induction of COX-mediated inflammation in chicken and frog embryos also caused abnormal NC migration and craniofacial development (Alvizi et al., 2023; Li et al., 2022). There appears to be a delicate balance between inflammatory and anti-inflammatory signaling that must be maintained for normal development to occur, but we still lack an understanding of the role of the COX signaling pathway within this balance.

Considering the types of developmental abnormalities described above, COX signaling appears to play a role in the normal development of NC and neural tube cells. NC cells are multipotent stem cells in the developing embryo that form within the neural tube, subsequently delaminate and undergo an epithelial-to-mesenchymal transition (EMT), and then migrate out into peripheral tissues where they give rise to a diverse array of derivatives (Achilleos and Trainor, 2012; Pla and Monsoro-Burq, 2018). NC derivatives range from melanocytes and cells of the peripheral and enteric nervous systems (PNS and ENS) to adipocytes, and craniofacial bone and cartilage (Achilleos and Trainor, 2012; Pla and Monsoro-Burq, 2018). Consequently, a diverse range of developmental anomalies can be attributed to abnormal NC cell development — termed ‘neurocristopathies’— including cleft palate, heart defects, and enteric neuropathies (Pilon, 2021).

To undergo EMT, migrate, and differentiate, NC cells alter their gene expression, cell-cell and cell-matrix adhesion, and cytoskeletal arrangements, and become migratory and invasive (Szabo and Mayor, 2018). Normal NC cell development, therefore, requires precise timing and a complex network of signaling pathways that drive the expression of key factors within their gene regulatory network (GRN). The complexity of these processes makes NC cells vulnerable to assaults including exposure to a variety of potential toxicants and teratogens that can disrupt these cellular processes (Bhagirath et al., 2021).

We investigated the effects of globally inhibiting COX signaling during development by NPX exposure in externally developing, vertebrate *Ambystoma mexicanum* (axolotl) embryos. Here, we confirm that embryonic expression of COX pathway genes is conserved across species. We additionally identify possible etiologies of NSAID-mediated developmental defects by characterizing the onset of developmental anomalies after NPX exposure and analyzing changes in the expression of key factors marking NC cell development. We show that exposing axolotl embryos to biologically relevant concentrations of NPX causes abnormal NC migration marked by a reduction in SOX9 protein expression. This reduced migration further manifests as stage-dependent changes in axial and craniofacial development marked by abnormal localization and patterning of collagen type II-positive (COL2A1+) craniofacial cartilage structures. Specifically, NPX exposure inhibits normal migration of NC cells fated to become craniofacial chondrocytes, leading to a hypoplastic, immature phenotype of Meckel’s cartilage. We also demonstrate that NPX exposure causes changes in the expression of peripheral and central nervous system (PNS and CNS) markers that may indicate an abnormal neurodevelopmental phenotype accompanying the changes in craniofacial chondrogenesis.

## II. Methods/Experimental Design

### Animal use and NPX exposures

Adult axolotls were housed and mated in accordance with the UC Davis Institutional Animal Care and Use Committee protocol #23047. Embryos were staged based on two factors: their developmental age and their gross morphology compared to control embryos from the same clutch (Adamson et al., 2022). The axolotls were cultured between 18°C - 22°C in Holtfreter’s (HF) solution (15 mM NaCl, 0.6 mM NaHCO3, 0.2 mM KCl, 0.2 mM CaCl2, and 0.2 mM MgSO4•7H20) for approximately 72 hours to reach stage 19 (early tailbud stage). Embryos were then either maintained in 100% HF solution or were exposed to various biologically relevant concentrations (5, 10, or 20 µg/mL) (Siu et al., 2002) of NPX (C14H14O3) dissolved in HF solution. Solutions were replaced once daily until the embryos reached the stage of interest for collection.

Embryos remained in their jelly coats during exposure and incubation. Embryos were cultured at room temperature (RT, approximately 21°C – 22°C) and were collected for immunohistochemistry (IHC) at one of three stages relating to NC cell development: stage 28 (specification and early migration), stage 36 (migration), and stage 45 (differentiation). Control embryos were monitored to determine the time it took to reach the developmental stages of interest and these timepoints were used to track developmental progress of the treated embryos. The development of key morphological characteristics such as the eye, gills, dorsal fin, and the degree of axial elongation were also weighed alongside developmental age to determine the stage of treated embryos. Once the embryos reached stages of interest, they were manually dejellied and dechorionated with forceps prior to fixation of the embryos in 4% paraformaldehyde (PFA).

### Western blot

Western blot (WB) was performed as previously described with some minor modifications (Chacon and Rogers, 2019). Embryo lysate was isolated from axolotl embryos cultured in 100% HF. We collected 10 embryos per stage from stages 26-28, 36-38, and 45 for analysis. Lysate was isolated using lysis buffer (50 mM Tris-HCL pH 7.4 with 150 mM NaCl plus 1.0% NP-40) and EDTA-free protease inhibitor (Roche complete, # 11697498001). Gel electrophoresis was performed on precast 4–12% bis-tris gel (Invitrogen, #NP0322BOX) for 1 hour at 40V, then 3 hours at 100V. The gel was transferred to a nitrocellulose membrane using the Invitrogen iBlot 2 Dry Blotting System. Nitrocellulose membranes were blocked and incubated in a milk blocking buffer solution of 5% milk protein in 1x TBS (500 mM Tris-HCl pH 7.4, 1.5 M NaCl, and 10 mM CaCl2) containing 0.1% Triton X-100 (TBST + Ca^2+^). Protein ladders used were: PageRuler Plus Prestained Protein Ladder, 10 to 250 kDa (Thermo Scientific) and MagicMark XP Western Protein Standard, 20 to 220 kDa (Invitrogen). No-Stain labelling (Invitrogen, #A44449) was performed before total protein imaging. The blots were then washed with ultrapure water and incubated in primary antibody (see Table 1) in the 5% milk protein + TBST + Ca^2+^ blocking solution overnight at 4°C. The blots were washed 3 times in 1x TBST + Ca^2+^ for 15 minutes per wash, then incubated in secondary antibody (Prometheus Protein Biology Products, #20-303) and visualized using an enhanced chemiluminescence (ECL) kit (GE Healthcare Lifesciences, #RPN2232).

**Table 1.**
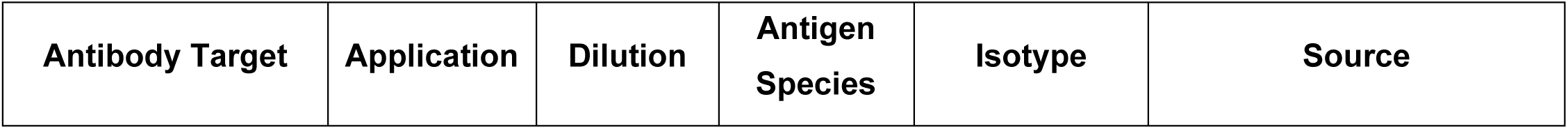
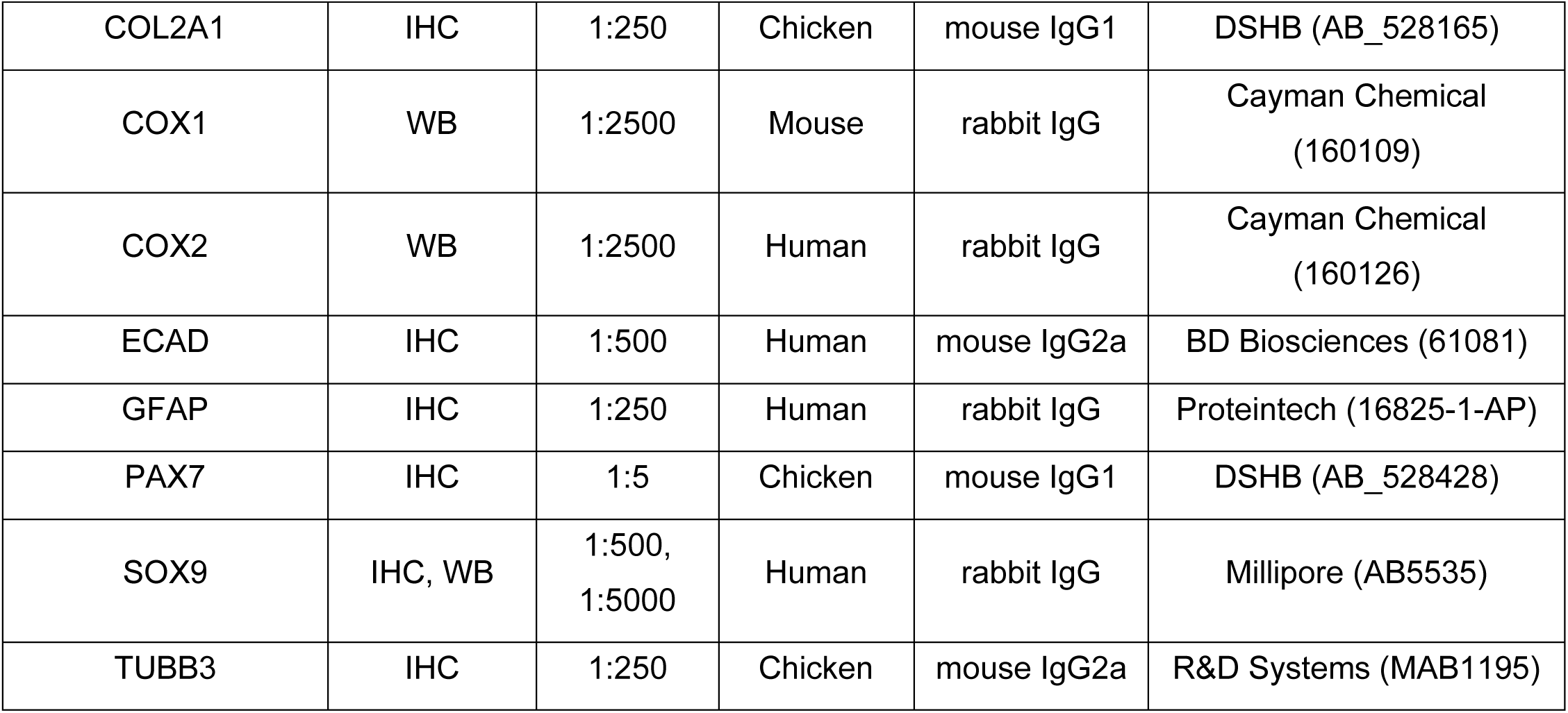
Primary antibodies used for IHC and WB.

Images were obtained using the Bio-Rad ChemiDoc MP Imaging System and image analysis was performed using ImageLab.

### Immunohistochemistry

IHC was performed as previously described with modifications (Monroy et al., 2022). Briefly, embryos were fixed in 4% PFA in phosphate buffer for 1 hour then quickly washed three times in 1x TBS (500 mM Tris-HCl pH 7.4, 1.5 M NaCl, and 10 mM CaCl2) containing 0.1% Triton X-100 (TBST + Ca^2+^) to remove remaining PFA. They were then incubated in a blocking buffer solution (10% donkey serum in 1x TBST + Ca^2+^) overnight at 4°C. Table 1 lists the primary antibodies used for IHC in this study. Antibodies were diluted (1:2-1:500) in blocking buffer with optimized dilutions for each antibody (Table 1). Embryos were incubated in primary antibodies and 4’,6-diamidino-2-phenylindole (DAPI) stain at 4°C for 48 hours. Next, embryos were washed in TBST buffer and secondary Alexa Fluor antibodies were added. Embryos were kept in the secondary antibody mixes overnight at 4°C, post-fixed in 4% PFA for 1 hour at RT, then washed in 1x TBST + Ca^2+^ six times for 15 minutes each wash prior to whole mount imaging.

### Fluorescence microscopy

Embryos were imaged in whole mount and after cryosectioning using the Zeiss Axio Imager.M2 microscope. Wholemount stage 28 and 36 embryos were imaged at 10X magnification, focused on the cranial region of the embryo while stage 45 embryos were imaged at 5X magnification. Embryos were washed in sucrose and embedded in gelatin for cryosectioning (see below). They were then cryosectioned at 16-25 µm thick sections. Cryosections were imaged using the ZEISS Apotome.2 at 10X-63X magnification. Images were obtained using ZEN Blue 3.0 optical processing software.

### Embedding and cryosectioning

Embryos were incubated in 5% sucrose at 4°C overnight, and then in 15% sucrose at 4°C overnight, then in 10% gelatin overnight at 37°C. Embryos were next embedded in 10% gelatin, flash-frozen in liquid nitrogen, and stored at -80° C until cryosectioning. Embryos were cryosectioned with a Microm NX70 cryostat.

### Image analysis

Bright field images were evaluated to determine changes in gross morphology of the developing embryos. Head length, dorsal fin height, eye diameter, and pre-optic length were all measured using Adobe Illustrator. For these measurements, line segments were converted to µm using in-image scale bars and the length was compared between treatments. Cell counts for SOX9 and PAX7+ cells were performed using the Adobe Photoshop counter tool as previously described by our lab (Monroy et al., 2022). Dorsoventral (DV) displacement was calculated as a measurement of cell migration; the methods for these calculations were adapted from a previously described statistical shape analysis technique (Sobhani et al., 2021). For DV displacement, Illustrator was used to draw three lines overlying the transverse section of the embryo: the first was drawn from the dorsal-most point on the section down to the ventral-most point (the total length of the DV axis); the second was drawn from the same starting point until the furthest migrated SOX9 or PAX7+ cell in the section; the last was drawn starting from that same cell and ending at the ventral point of the DV axis. The DV displacement was calculated using these measurements and trigonometric equations to represent the migration distance of cranial NC cells in each transverse section to account for differences in embryo sizes. Additional details are provided in Supplemental Figure 8.

Morphometric analysis of the developing cartilage was performed using Photoshop and FIJI (ImageJ). The total area and circularity of each cross-section through Meckel’s cartilage was measured using the built-in image analysis tools in Photoshop. The lasso tool was used to highlight Meckel’s cartilage based on the COL2A1 positivity, and then the area and circularity of this selection were analyzed using the image analysis and measurement tools. Cell density was calculated using the measured total area of the cross-sections and cell counts of the DAPI+ cells within the COL2A1+ region which were obtained using the lasso selection tool and counter tool, respectively. Then, using FIJI, the cross sectioned images of Meckel’s cartilage were thresholded into two images for separate analysis: (1) perichondral COL2A1+ band and (2) intra-articular COL2A1+ bundles. For the collagen band, the ImageJ plug-in *MorphJ* was utilized to determine local thickness and area of staining. For the intra-articular bundles, thresholded images were watershed and internal function *“analyze particles”* was utilized to count and determine the area of each bundle.

GraphPad Prism 10 was used to generate violin plots for the cell count, DV displacement, and morphometric data obtained from bright field and wide field fluorescence microscopy image analysis. Statistical analysis using nonparametric, unpaired Mann-Whitney tests was performed using Prism. The relevant sample sizes used for quantitative analyses, as well as the total sample size for each protein of interest in our IHC experiments, are listed in the individual figure captions. For each embryo, multiple transverse sections were analyzed. Z-scores for Figure 5 describe the axial level at which the image of the transverse section was taken and were determined by identifying the presence of ocular structures within the section included the developing cornea, retina, and lens. Based on the prescence or absence of these structrues, the following z-scores were defined: pre-lens, lens, post-lens, and post-optic.

### Single-cell sequencing analysis

Publicly available embryonic single cell sequencing data sets were obtained and variably processed according to the organism corresponding to each dataset. Detailed information is described in Supplementary Table 1. All plots were created using built-in Seurat functions (Hafemeister and Satija, 2019) or ggplot (Wickham, 2016) and were organized in BioRender.

## III. Results

### Analysis of COX signaling pathway transcript expression during embryogenesis across species

Previous studies using zebrafish embryos showed that members of the COX pathway are expressed during early embryonic development (Cha et al., 2005; Cha et al., 2006b). Additionally, the COX signaling pathway is proposed to have a role in several key developmental processes including gastrulation based on COX activity and the effects of NSAIDs during early development (Cha et al., 2006a; Grosser et al., 2002; Ishikawa et al., 2007). To confirm the conservation, and relevance, of the COX pathway factors during vertebrate embryogenesis, we utilized publicly available whole embryo single cell RNA-sequencing datasets from zebrafish (Farnsworth et al., 2020; Farrell et al., 2018; Raj et al., 2020; Wagner et al., 2018), frog (Petrova et al., 2024), mouse (Pijuan-Sala et al., 2019), and macaque (Zhai et al., 2022). We selected datasets which incorporated all three germ layers and were staged around early organogenesis. Specifically, the zebrafish dataset represented 12 hours post fertilization through 2 days post fertilization, the African clawed frog was from stage 18 through stage 22, the house mouse was from mixed late gastrulation through embryonic day 8.5, and the cynomolgus macaque dataset ranged from embryonic day 20 through 29.

Within the anamniotes, cells grouped into 25 clusters within the zebrafish and 25 clusters in the frog (Figure 1A, B). We noted a distinct expression of major cyclooxygenase pathway enzyme genes, *ptgs1* (*COX- 1*), *ptgs2* (*COX-2*) and *ptges*, in the ectodermal lineage clusters such as the epidermis and neural crest (Figure 1C-N). In addition, there was a notable expression within the mesoderm lineage (Figure 1C-N). When comparing expression of COX pathway genes with amniotes, we noted high variability in expression patterns of these 3 genes across the 36 mouse and 37 macaque cell types (Figure 2A-N). Most notably, *PTGS1* and *PTGS2* were largely restricted to the hemato-endothelial types in both species (Figure 2C-F, 2K-N) and *PTGES* was broadly expressed in the mouse but absent in the macaque (Figure 2G, H, M, N). In the macaque, we noted a low but distinct *PTGS1* and *PTGS2* expression in the ectodermal derivatives such as brain, spinal cord and neural crest (Figures 2I-L). Other COX pathway elements were also identified sporadically but minimally across all species. In particular, *PTGES3* was found to be ubiquitously expressed across most cell types in all species (Supplemental Figures 1-4).

**Figure 1.**
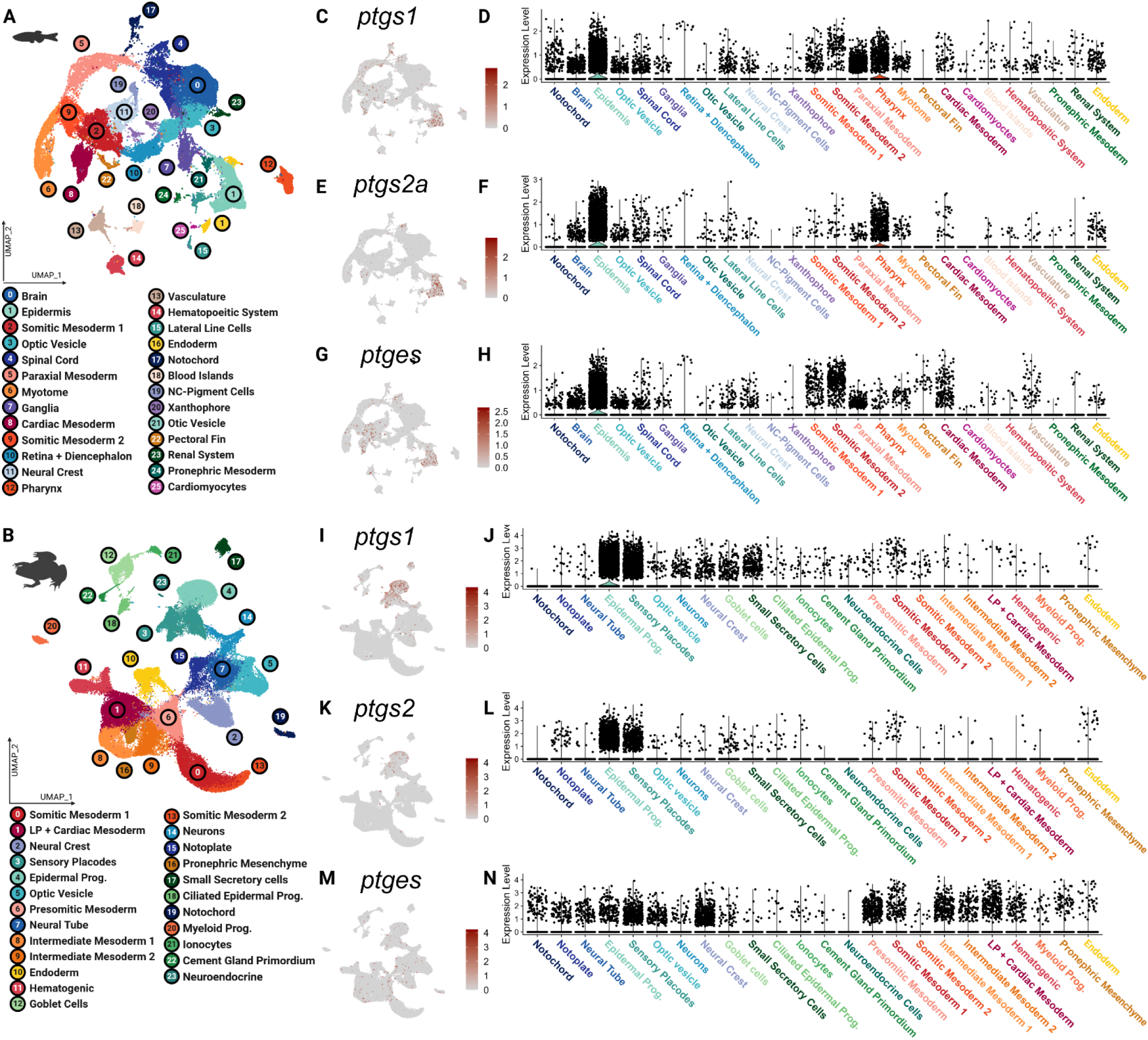
Cyclooxygenase pathway is active in multiple embryonic cell types across anamniotes. Analysis was performed of publicly available scRNA-seq data for whole embryos, staged around early organogenesis for the zebrafish (*Danio rerio*) and African clawed frog (*Xenopus laevis*). (A, B) UMAP demonstrating clustering results of whole embryos from the anamniotes. Feature UMAPs and violin plots demonstrating expression of select major enzymes of the cyclooxygenase pathway— *COX-1*/*ptgs1* (C, D, I, J), *COX-2*/*ptgs2* (E, F, K, L) and *ptges* (G, H, M, N)— across the major cell types identified by species. NC, neural crest; LP, lateral plate; Prog., progenitor.

**Figure 2.**
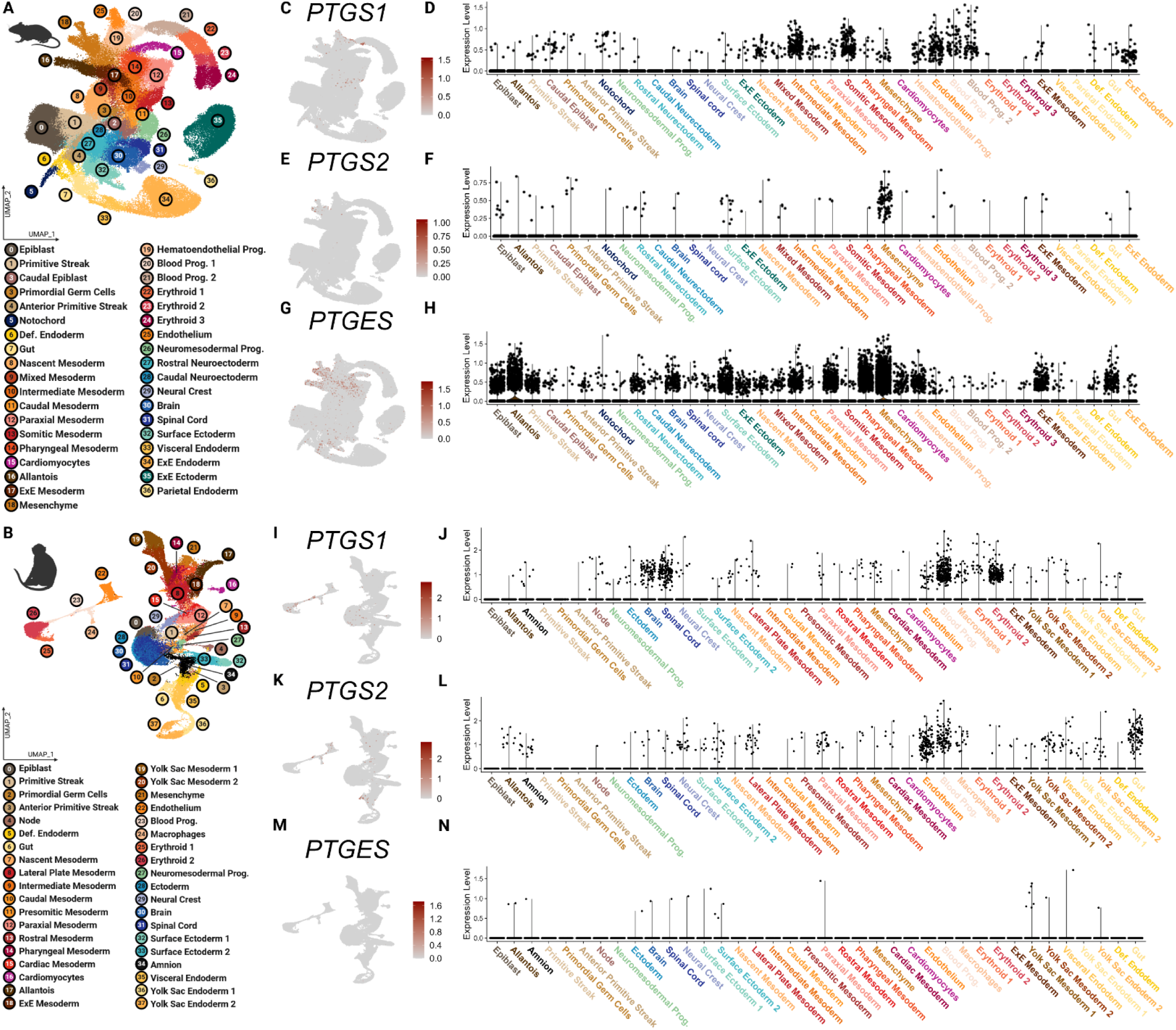
Cyclooxygenase pathway is active but more restricted in embryonic cell types across amniotes. Analysis was performed of publicly available scRNA-seq data for whole embryos, staged around early organogenesis for the house mouse (*Mus musculus*) and crab-eating macaque (*Macaca fascicularis*). (A, B) UMAP demonstrating clustering results of whole embryos from the amniotes. Feature UMAPs and violin plots demonstrating expression of select major enzymes of the cyclooxygenase pathway— COX-1/*PTGS1* (C, D, I, J), COX-2/*PTGS2* (E, F, K, L) and *PTGES* (G, H, M, N)— across the major cell types identified by species. Prog., progenitor; Def., definitive; ExE, extra embryonic.

To confirm expression of COX signaling proteins during axolotl embryogenesis we performed western blot and identified expression of COX pathway proteins COX-1 and COX-2, across several stages of axolotl development spanning from NC specification (stage 26-28) through early differentiation (stage 45) of the NC into its various derivatives (Figure 3A, B). We confirmed that these proteins are co-expressed at stages where NC cells are developing using an antibody against SOX9 (Figure 3C). Quantification showed that both COX-1 and COX-2 expression is higher during NC formation, migration, and early differentiation stages than in later stage 45+ tadpoles.

**Figure 3.**
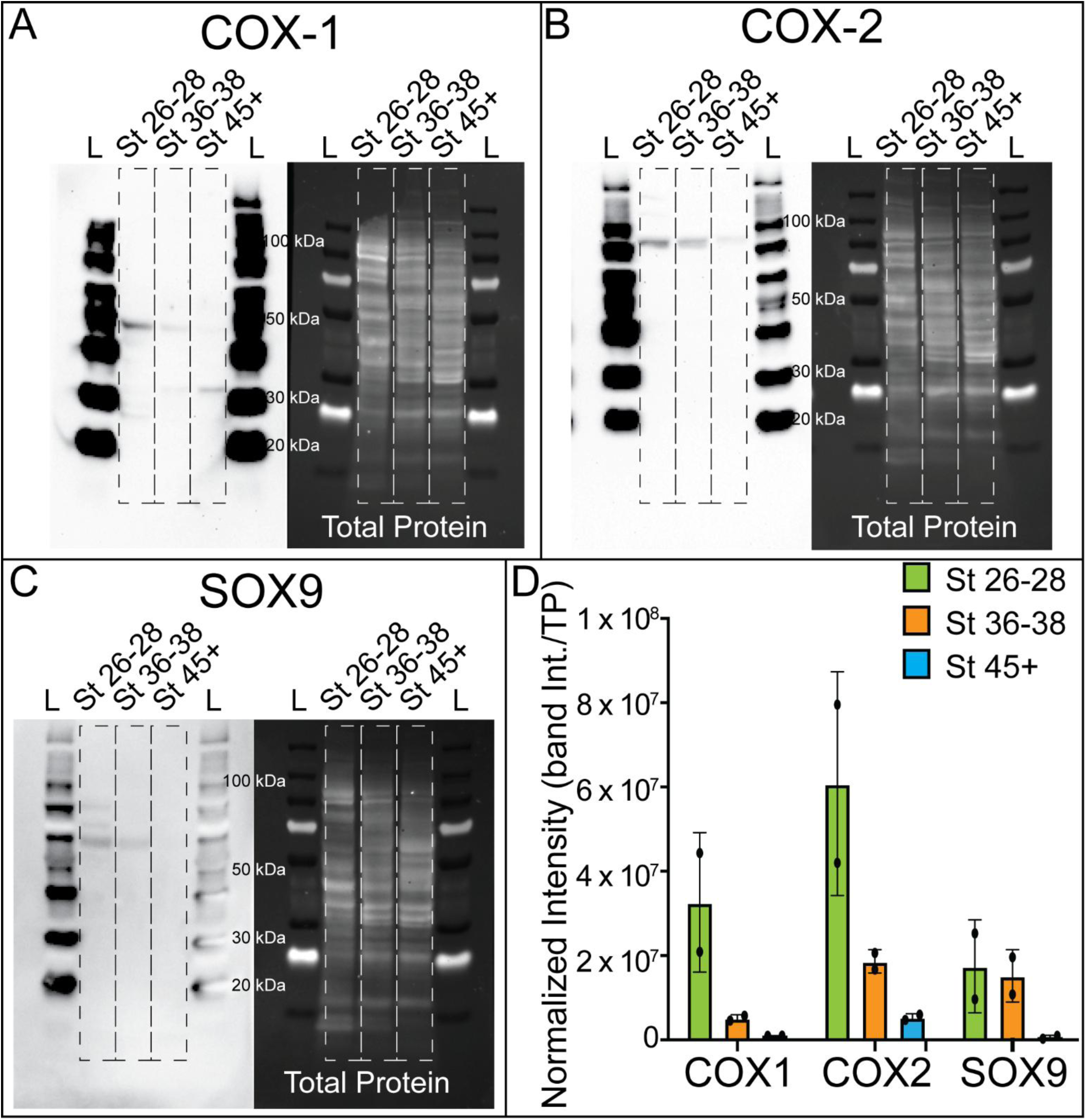
COX pathway proteins are expressed across stages of NC development in the axolotl. (A-C) Western blot showing expression of COX pathway proteins (A) COX-1, (B) COX-2, and NC marker, (C) SOX9, from protein lysate collected from stage 26-28, 36-38, and 45+ axolotl embryos. Chemiluminescent blots (left) and total protein images (right) shown. (D) Quantification of the band intensity of the WB in panel A normalized to the total protein per lane. ImageLab software (BioRad) used for quantification of total protein and band intensity.

### NPX exposure causes morphological defects

Prior research showed that COX inhibition through COX-2 selective and non-selective NSAID exposure does in fact cause significant developmental defects in NC-derived structures including cardiac malformations, abnormal innervation of the PNS and ENS, and craniofacial abnormalities (Hultzsch et al., 2021; Nakhai-Pour and Berard, 2008; Schill et al., 2016; Yoon et al., 2018). Since prior work identified defective embryonic development with ibuprofen and celecoxib exposures, we hypothesized that there would be observable morphological changes in axolotl embryo development following exposure to the non-selective NSAID, NPX. To test this hypothesis, we exposed axolotl embryos to varying biologically-relevant concentrations of NPX (Siu et al., 2002) and performed morphological analysis to evaluate the incidence and severity of developmental anomalies. Embryos were first exposed to NPX at late neurula and early tailbud stages (∼stage 20-24) to focus on NC formation and migration stages and to avoid germ layer malformation during gastrulation. We found that inhibition of COX signaling through NPX exposure during these stages caused a reduction in the survival of axolotl embryos in a dose-dependent manner (Supplemental Figure 5). In stage 28 axolotl embryos (tailbud), there were no overt effects on morphological development observed independent of the degree of COX inhibition via NPX exposure (Figure 4A-D). However, in stage 45 axolotl tadpoles, we identified obvious defects in growth and morphology (Figure 4E-H). Specifically, compared to control tadpoles, all those exposed to any concentration of NPX had defects in head length (Figure 4E-H, I), dorsal fin height (Figure 4E-H, J) eye diameter (Figure 4E-H, K), and pre-optic length (Figure 4E-H, L). Considering the low rates of survival and concentration-dependent scale of defects, it is clear that inhibition of COX signaling broadly alters axolotl growth and development.

**Figure 4.**
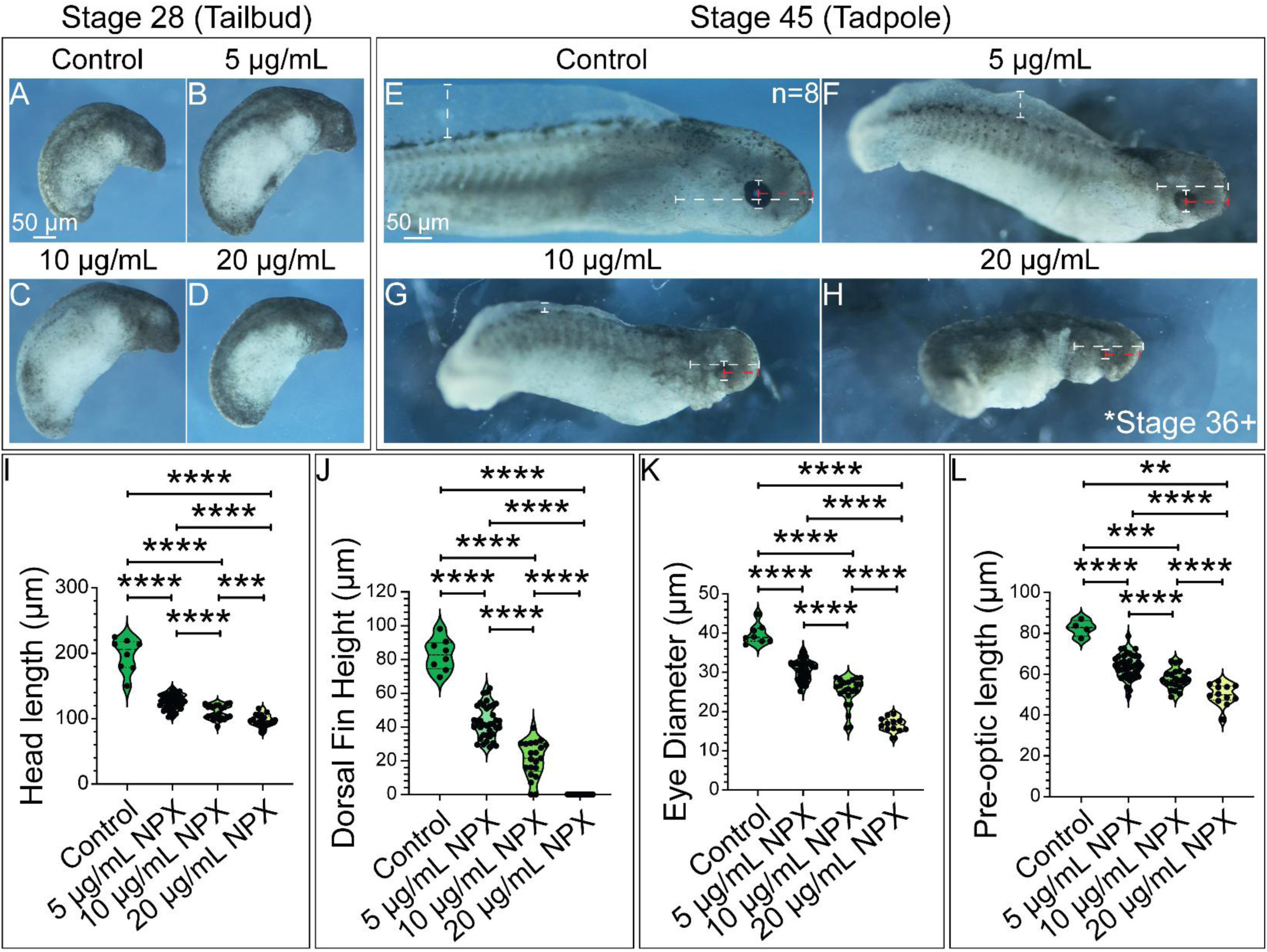
The degree of NPX exposure corresponds to the severity of morphological effects observed in late-stage axolotl embryos. (A-D) Whole mount stage 28 embryos with anterior to the top right. (E-H) Whole mount stage 45 embryos with head to right. Embryos exposed to 20 µg/mL NPX developed to stage 36 collection timepoint but did not reach stage 45. They are included here to compare morphology in late-stage embryos and are labelled as Stage 36+ to clarify this inconsistency. Dashed bars demonstrate how morphologic measurements were performed. (I-L) Graphs showing quantification of head length, dorsal fin height, eye diameter, and pre-optic length in stage 45 embryos. Sample sizes for morphologic analysis were n=8, n=40, n=22, and n=19 for the control, 5 µg/mL NPX, 10 µg/mL NPX, and 20 µg/mL NPX treatment groups, respectively. Embryos that did not have grossly appreciable development of their eyes or gills were excluded from the eye diameter, pre-optic length, and head length measurements. Non-parametric, unpaired Mann-Whitney t-tests were performed to analyze these results. p-value= *, **, ***, **** = p-value < 0.05, < 0.01, < 0.001, and <0.0001, respectively.

Non-selective NSAIDs such as ibuprofen and indomethacin have been linked to various NC-specific defects; however, it is unclear if the observed teratogenic effects of NSAIDs are COX isoform-specific. To address this gap in knowledge, we performed additional exposure experiments using another non-selective NSAID, ibuprofen, and the COX-2-selective NSAID, celecoxib. When exposed to high concentrations of ibuprofen, axolotl embryos developed similar axial and craniofacial defects as those seen in the 20 µg/mL NPX-treated embryos (Supplemental Figure 6A-D compared to Figure 4E-H). In contrast, following celecoxib treatment, IHC against COL2A1 for the cranial cartilage structures revealed no overt craniofacial phenotypes when compared to control embryos (Supplemental Figure 6G, J compared to E, H). The distinct phenotypes observed following exposure to non-selective NSAIDs versus COX-2-selective NSAIDs suggest that the activity of COX-1 may be essential for the normal development of NC-derived tissues. As such, we used NPX for all subsequent exposure studies and measured its effect on expression of key regulators of NC development that could constitute a molecular mechanism for the development of the observed defects.

### COX inhibition impedes NC migration

Previous research demonstrated that ibuprofen exposure affects the migration and bowel colonization of NC-derived cells (Schill et al., 2016), and paired with the gross morphological changes observed in our embryos following NPX exposure (Figure 4), we wanted to determine if COX signaling is necessary for early NC development. To identify if COX signaling is necessary for NC cell specification and migration, we exposed embryos at late neurula and early tailbud stage to various concentrations of NPX and then used IHC to evaluate changes in expression of two key transcription factors that regulate early NC cell development, SOX9 and PAX7. We identified that although the tailbud embryos appeared morphologically normal in wholemount (Figure 4A-D), compared to control embryos, COX inhibition via NPX exposure at 5 µg/mL, 10 µg/mL, and 20 µg/mL reduced the numbers of SOX9+ and PAX7+ cranial NC cells (Figure 5A-L, Q). In addition to reducing the number of NC cells, NPX exposure reduced the migration distance of these cells (Figure 5R, Supplemental Figure 8). When we quantified dorsal-ventral (DV) displacement, we determined that NPX exposure significantly reduced the migration distance of SOX9+ NC cells at 20 µg/mL NPX concentration compared to control embryos (Figure 5E-H, M-P, R). While SOX9 marks the cranial NC lineage that is fated to become craniofacial cartilage, PAX7 is necessary for NC specification and induction of downstream effectors in the NC GRN upstream of SOX9 (Basch et al., 2006). We also identified significant changes in the ability of PAX7+ cells to migrate away from the dorsal neural tube following NPX exposure, and DV displacement was significantly reduced in the 10 µg/mL and 20 µg/mL groups compared to control embryos (Figure 5I-L, M-P, R). These results suggest that COX signaling may be necessary for NC specification and migration.

**Figure 5.**
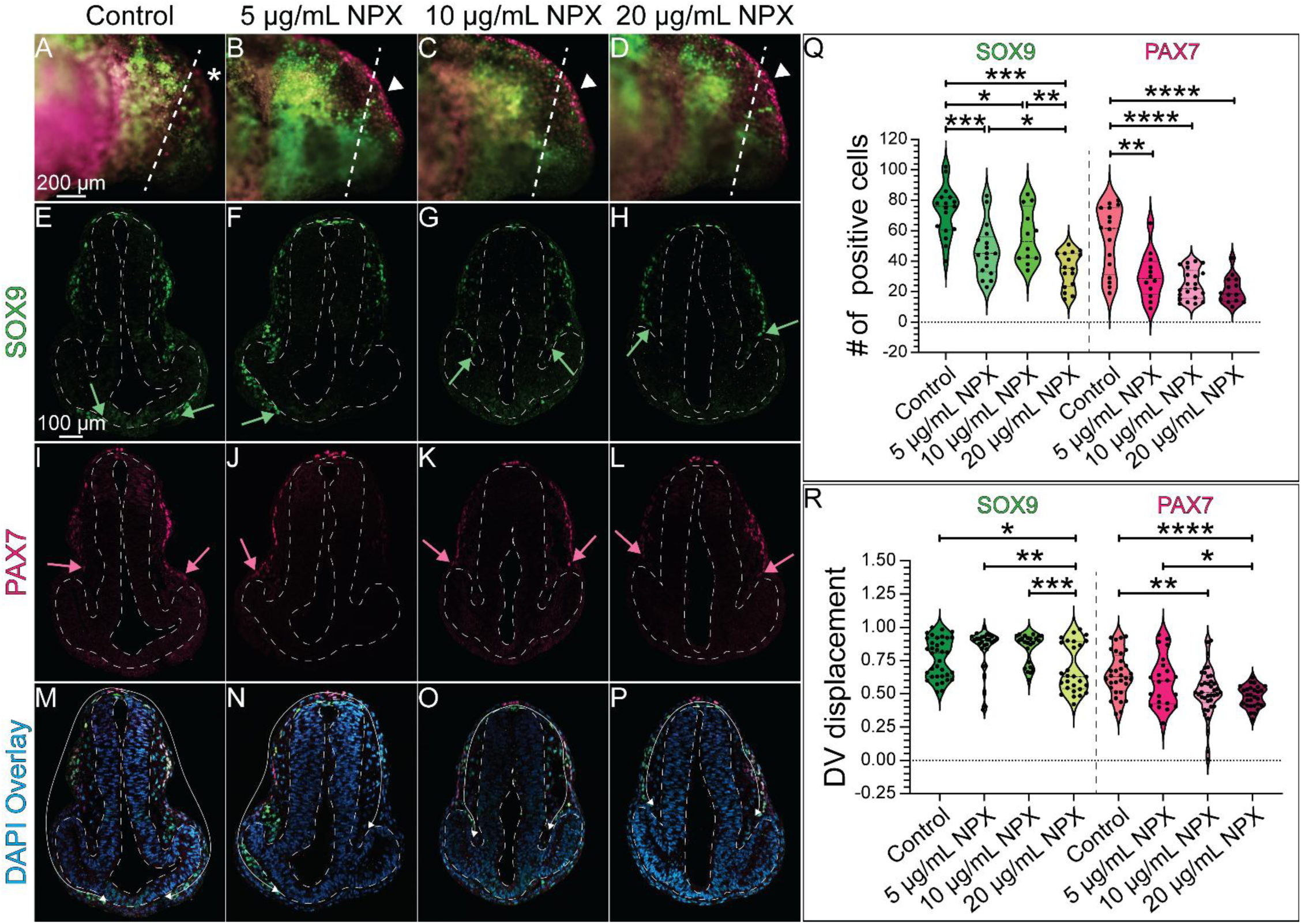
NPX exposure reduces the migration of SOX9+ and PAC7+ cells into the developing cranial region. IHC for SOX9 and PAX7 in (A-D) whole mount and (E-P) transverse sections of stage 28 axolotl embryos. (A-D) Dashed lines show the axial level in the head that corresponds to the transverse sections. Arrowheads highlight cranial PAX7 expression which appears abnormally localizes in (B-D) treated embryos compared to PAX7 expression in the (A) control embryos marked by the asterisk. (E-P) Dashed lines outline the neural tube and developing eyes in transverse section. (E-H) Green arrows indicate the extent of migration of SOX9+ NC cells in the cranial region. (I-L) Pink arrows indicate the extent of migration of PAX7+ NC cells. (M-P) Overlay of SOX9 and PAX7 with DAPI. The white arrows show migratory path of NC cells into the ventral face. (Q) Quantification of the number of SOX9+ or PAX7+ cranial NC cells. (R) Quantification of the migration distance of SOX9+ or PAX7+ NC cells, measured as DV displacement, in transverse section. For SOX9 analysis, the following sample sizes were used: n=6, n=5, n=5, and n=6 embryos for the control, 5 µg/mL NPX, 10 µg/mL NPX, and 20 µg/mL NPX groups, respectively, with an average of 3 transverse sections analyzed per embryo. Similarly, for PAX7 analysis, the sample sizes were n=5, n=5, n=6, and n=6 embryos for the respective treatments, and an average of 3 transverse sections were analyzed per embryo. The number of data points in the violoin plots reflects the total number of transverse sections analyzed per marker rather than representing a single average value across sections for each individual embryo (biological replicate). The number of embryos per treatment group that recapitulate the phenotype shown in this figure are n=25, n=21, n=29, and n=16 for the respective treatments. Non-parametric, unpaired Mann-Whitney t-tests were performed to analyze these results. p-value= *, **, ***, **** = p-value < 0.05, < 0.01, < 0.001, and <0.0001, respectively.

### Global COX inhibition by NPX alters cranial cartilage formation

Based on the reduction in the number and migratory capacity of SOX9+ and PAX7+ NC cells following NPX exposure, we hypothesized that there may be downstream effects on NC cell differentiation into cranial chondrocytes. We exposed embryos to the same concentrations of NPX, performed IHC for COL2A1, and then evaluated changes in the development of cranial cartilage structures using morphometric analysis (Figure 6).

**Figure 6.**
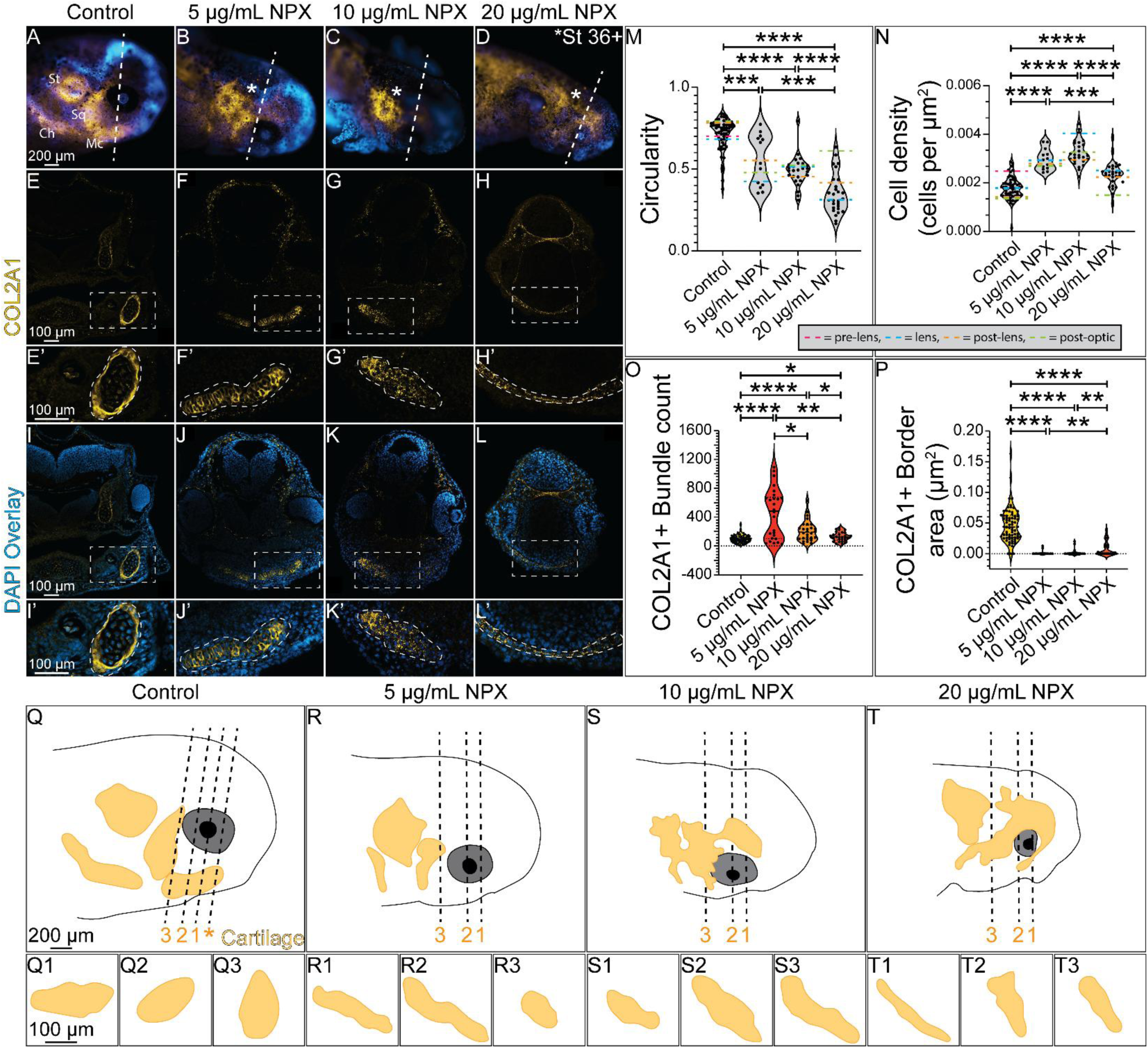
COX inhibition causes abnormal formation of cranial cartilage structures. IHC for COL2A1 in (A-D) wholemount and (E-L) transverse sections from stage 45 axolotls. (A-D) Dashed line indicates axial level of sections in (E-L). Asterisks mark abnormal COL2A1 localization. (E-L) Dashed boxes indicate region of zoom in. (E’-L’) Enlarged images of Meckel’s cartilage show abnormal tissue morphology and COL2A1 localization following NPX exposure. Dashed outlines highlight the developing chondrocytes within Meckel’s cartilage expressing COL2A1. (D, H, L) Embryos exposed to 20 µg/mL NPX developed to stage 36 collection timepoint but did not reach stage 45. They are included here to compare cartilage morphology in late-stage embryos and are labelled as Stage 36+ to clarify this inconsistency. (M-P) Quantification of circularity and cell density of Meckel’s cartilage with the number of COL2A1+ bundle structures within Meckel’s cartilage and the total area of the COL2A1+ band surrounding Meckel’s cartilage in cross section. Pink, green, orange, and blue dashed lines on the violin plots in M-N indicate the average circularity or cell density values at each axial level (pre-lens= pink, lens= blue, post-lens= orange, and post-optic= green) across the 4 treatment groups. For analysis of COL2A1, the sample sizes analyzed were n=6, n=5, n=4, and n=4 for the respective treatment groups. The total number of embryos per treatment group that express similar phenotypes are as follows: n=18, n=29, n=21, and n=9, respectively. An average of 4 transverse sections were analyzed per embryo. The number of data points in the violoin plots reflects the total number of transverse sections analyzed per marker rather than representing a single average value across sections for each individual embryo (biological replicate). (Q-T) Drawings of wholemount stage 45 axolotl embryos in A-D with the developing COL2A1+ cranial cartilages drawn in the yellow structures. Dashed lines indicate the axial levels that were analyzed in the circularity and cell density analysis of Meckel’s cartilage in M-N (asterisk= pre-lens, 1 = lens, 2 = post-lens, and 3 = post-optic). (Q1-Q3, R1-R3, S1-S3, and T1-T3) Drawings of transverse sections through Meckel’s cartilage at each axial level of interest across treatment groups. Ch= ceratohyal cartilage, Mc= Meckel’s cartilage, Sq= squamate cartilage, St= supratemporal cartilage. Non-parametric, unpaired Mann-Whitney t-tests were performed to analyze these results. p-value= *, **, ***, **** = p-value < 0.05, < 0.01, < 0.001, and <0.0001, respectively.

Alterations in COL2A1 expression were apparent in wholemount stage 45 tadpoles (compare Figure 6A to 6B- D and 6Q to 6R-T). Analysis of COL2A1 in transverse cryosections revealed that in comparison to control tadpoles, which had normal mandibular cartilage (Meckel’s cartilage) morphology, embryos exposed to all concentrations of NPX had abnormal development of Meckel’s cartilage demonstrated by abnormal localization of COL2A1 (Figure 6F-H, J-L). Maturing chondrocytes express high levels of collagen types II and IV and proteoglycans like aggrecan, which help draw water into the cartilage tissue, giving the tissue its shape (Zheng et al., 2024). We indirectly evaluated chondrocyte maturation and cartilage tissue morphogenesis by quantifying the circularity and the cell density of Meckel’s cartilage in cross-section. We further quantified the fluorescence COL2A1signal (bundles) within each cross-section and the total area of the COL2A1+ border band (perichondrium) that surrounds Meckel’s cartilage in control embryos.

We identified abnormal Meckel’s cartilage morphology in treated embryos based on the reduction in cross-sectional circularity and elongation of Meckel’s cartilage in cross-section in NPX-exposed embryos compared to control embryos (Figure 6E’-H’, I’-L’, M). The increase in cell density within Meckel’s cartilage in treated embryos compared to control embryos (Figure 6I’-L’, N) suggests that the chondrocytes in the treated embryos are not developing normally, further indicating that exposure to NPX inhibits normal chondrocyte maturation and cartilage morphogenesis. Although all tadpoles expressed COL2A1 to some extent, the localization of the signal within the cross-sections of Meckel’s cartilage varied. In control embryos, there were fewer and smaller bundles of COL2A1+ clusters inside a discrete border of COL2A1-expressing cells around the outside of Meckel’s cartilage (Figure 6E’, I’, O-P). In contrast, in treated embryos there were no distinct COL2A1+ perichondrial border bands of tissue and there were increased COL2A1+ bundles than in control embryos (Figure 6F’-H’, O-P). Representative illustrations of COL2A1+ cranial cartilages in wholemount stage 45 axolotl embryos demonstrate the consequences of NPX exposure on patterns of COL2A1 expression in the developing head (Figure 6Q-T). Focusing then on cross-sections of Meckel’s cartilage across several axial levels within the head, the morphology of Meckel’s cartilage was irregular following NSAID exposure across different regions of Meckel’s cartilage (Figure 6Q1-Q3, R1-R3, S1-S3, and T1-T3). These results demonstrate that there are clear defects in NC-derived craniofacial cartilage associated with NPX exposure, suggesting that reduced NC cells from earlier stages do not recover.

### COX signaling inhibition alters ectodermal derivative formation

As we observed a reduction in the number of PAX7 and SOX9+ NC progenitors following NPX exposure, we hypothesized that other NC derivatives may also be affected by COX inhibition following NPX exposure. NC drivatives include pigment cells, glia, and sensory neurons of the PNS. The pigment cell types present in axolotls— including melanophores, xanthophores, and iridophores— are also NC-derived and their abundance and distribution in the skin of developing axolotls has been previously described (Frost et al., 1984). Previous work in zebrafish showed that PAX7 is an important regulator of melanophore and xanthophore differentiation (Nord et al., 2016), so we evaluated PAX7 expression in stage 45 embryos to identify possible effects of NPX exposure on NC-derived pigment cells. In whole mount images, we identified that the normal patterning of pigment cell progenitors marked by PAX7 expression in the cranial region (Figure 7A-A’, B-B’) is abnormal in the NPX-treated embryos (Figure 7D-D’, E-E’). In transverse sections from control embryos, many PAX7+ pigment cells were restricted to the region underlying the ECAD+ epidermis (Figure 7C, C’, pink arrows). In contrast, in NPX treated sections, the integrity of the Ecad+ cells appeared disorganized and localization of PAX7 was altered. Specifically, PAX7+ cells in the NPX-treated embryos colocalized in cells expressing ECAD (Figure 7F, F’, G). We evaluated expression of ECAD in transverse sections to identify changes in the formation and structure of epidermal sensory organs. These organs of the lateral line sensory system in axolotls are composed of NC-derived glial cells and placodally-derived sensory neurons, and they have well-characterized morphology in the axolotl that can be evaluated through histology (Northcutt, 1992; Northcutt et al., 1994). IHC for ECAD revealed that the complex architecture of sensory organs in the skin of control embryos (Figure 7C, C’) was completely lost following exposure to NPX (Figure 7F, F’, G). These data suggest that the development of multiple major lineages of cranial NC cells are affected by early exposure to NPX. In consideration with the data shown in previous figures, there is a strong correlation between exposure to NPX and abnormal patterning, differentiation, and morphology of all ectodermally derived tissue structures.

**Figure 7.**
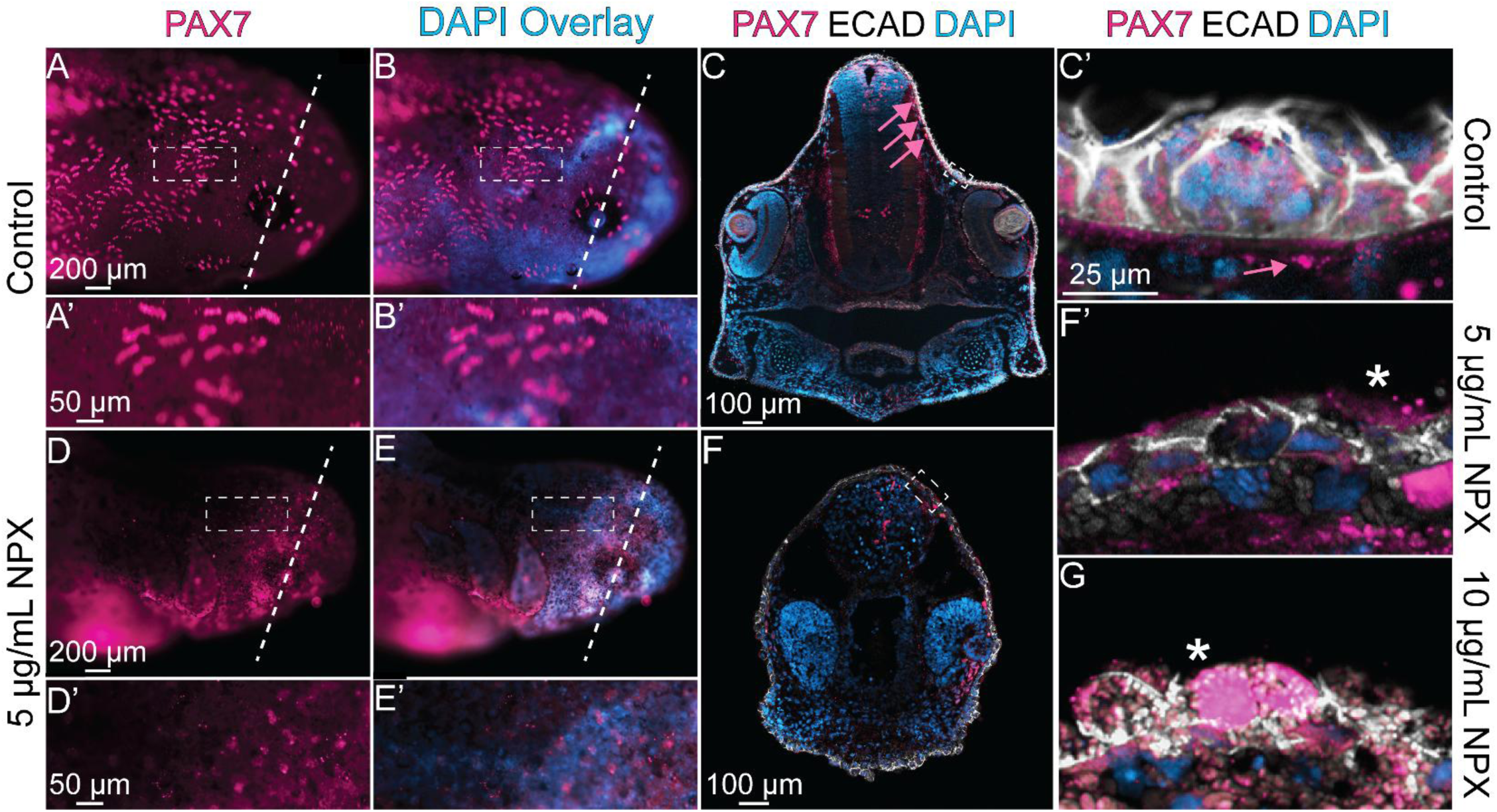
NPX alters PAX7 expression during the formation of epidermal sensory structures. IHC for PAX7 and ECAD in (A, B, D, and E) whole mount and (C, F) transverse sections. (A’, B’, D’, and E’) Zoom in from A, B, D, and E (dashed boxes). (A-E, A’-E’) The number of embryos that reflect the phenotype shown here are n=27 and n=14 for the control and 5 µg/mL NPX groups, respectively. (C) In control embryos, PAX7 and ECAD are localized to pigment cells and superficial sensory organs. Pink arrows highlight subepidermal localization of PAX7+ pigment cells. Dashed box identifies an ECAD+ sensory organ. (C’, F’) Zoom in from sections in C and F (dashed boxes). The pink arrows in panel C’ again highlights the position of PAX7 expression relative to the sensory structures in the skin. Four separate images were taken of the control embryo to fit the entire transverse section in the imaging field; these were then stitched together using the automated photomerge feature in Adobe Photoshop. (F’, G) Sensory organs from exposed embryos develop abnormally by stage 45. Embryos exposed to 20 µg/mL NPX developed to stage 36 collection timepoint but did not reach stage 45, and therefore, they are not included here. Asterisks indicate mislocalization of PAX7 expression.

Abnormal craniofacial development has an increased association with neurodevelopmental disorders (Tillman et al., 2018). Additionally, previous work identified that in embryos with abnormal NC cell development, there are often defects in other ectodermally-derived structures (Trainor, 2010). The development of NC and neural tube cells are tightly associated due to their physical proximity in the embryo, which means they share a very similar signaling landscape and receive similar mechanical cues while developing (Crane and Trainor, 2006). Prior work from our lab has also identified expression of COX pathway factors in the developing CNS of chicken embryos (Leathers et al., 2024a; Leathers et al., 2024b). Based on this information, we hypothesized that NPX exposure may have effects on CNS development in addition to the described effects on NC cells and NC-derived tissues. To test this hypothesis, we exposed embryos to NPX as in prior experiments and subsequently performed IHC for β-III tubulin (TUBB3) and glial fibrillary acidic protein (GFAP). At the stages assessed, TUBB3, which marks neuronal microtubules, is expressed in developing neurons in the PNS and CNS (Figure 8A, B, D) (Chacon and Rogers, 2019). Fluorescence intensity analyses on transverse cryosections from control and NPX-exposed embryos revealed that the signal was significantly reduced in embryos exposed to NPX at the 5 µg/mL and 10 µg/mL NPX concentration (compare Figure 8B, D, M to Figure 8F, H, J, L, M). Similarly, the GFAP fluorescence signal in the forming CNS was also reduced in 10 µg/mL NPX-exposed embryos compared to control embryos (compare Figure 8C, D, N to Figure 8K, L, N). These results, along with those shown in previous figures, suggest that there is an overall decline in the development of both NC and CNS derivatives after NPX exposure.

**Figure 8.**
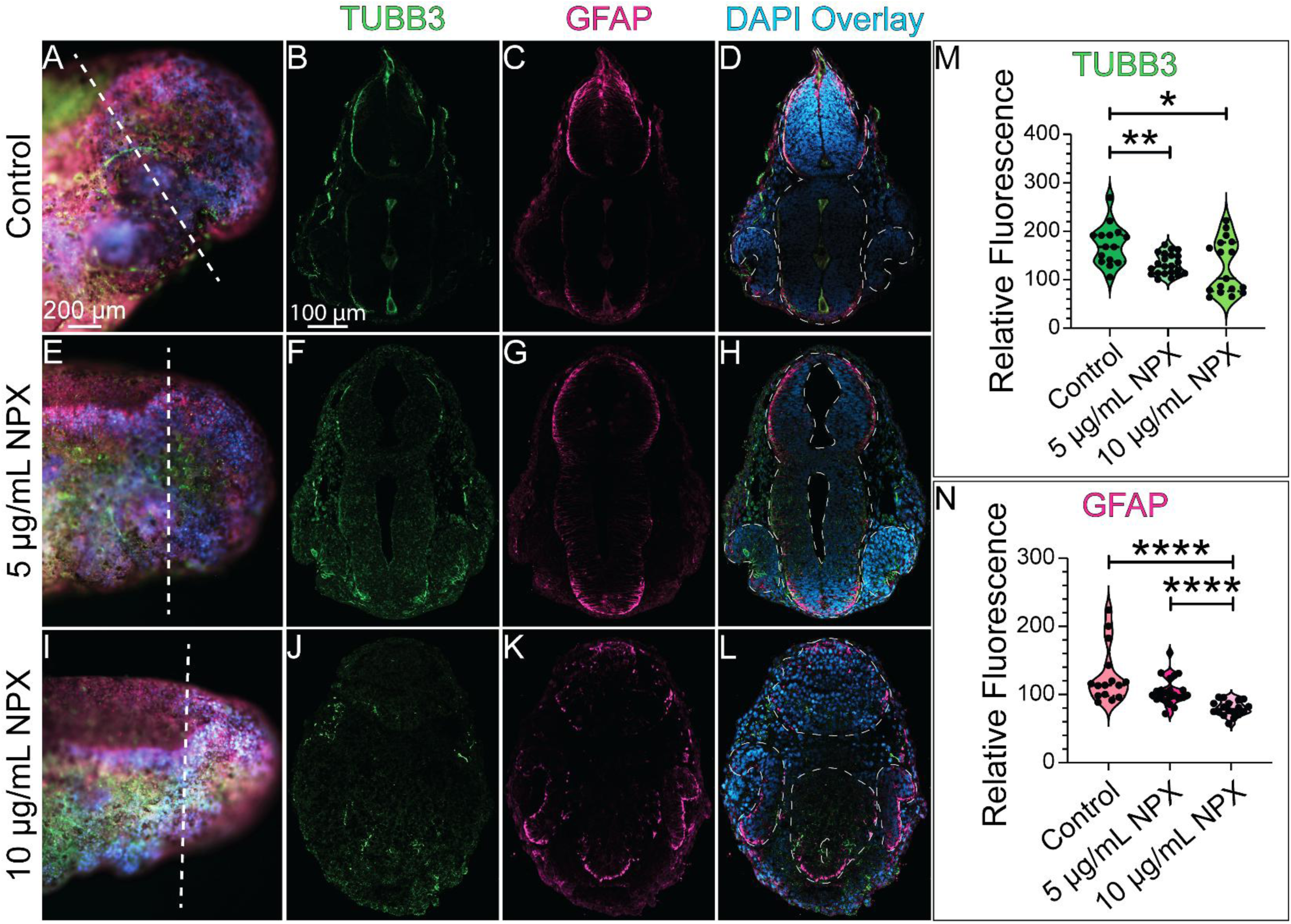
The expression of key neurodevelopmental markers is reduced following NPX exposure. IHC for TUBB3 and GFAP shown in (A, E, I) wholemount and (B-D, F-H, and J-L) transverse sections in (A-D) control embryos and (E-L) NPX exposed embryos. (A, E, I) Dashed lines show the axial level of transverse sections. (B-D, F-H, and J-L) Compared to control, NPX exposed embryos show reduced expression of TUBB3 (neurons) and GFAP (astrocytes/glia) in the neural tube. (D, H, L) Dashed outines highlight the developing neural tube and eyes. (M, N) Quantification of TUBB3 and GFAP fluorescence intensity in section. The sample sizes used for quantification of both TUBB3 and GFAP expression were n=5, n=5 and n=5 embryos for the control, 5 µg/mL, and 10 µg/mL NPX groups. An average of 4 transverse sections per embryo were quantified for both TUBB3 or GFAP expression. The number of data points in the violoin plots reflects the total number of transverse sections analyzed per marker rather than representing a single average value across sections for each individual embryo (biological replicate). The total number of embryos per treatment group expressing the phenotype shown in this figure is n=18, n=15, and n=26 for the respective treatments. Embryos exposed to 20 µg/mL NPX developed to stage 36 collection timepoint but did not reach stage 45, and therefore, they are not included here. Non-parametric, unpaired Mann-Whitney t-tests were performed to analyze these results. p- value= *, **, ***, **** = p-value < 0.05, < 0.01, < 0.001, and <0.0001, respectively.

## IV. Discussion

### Summary

Prior research demonstrated that exposure to NSAIDs during early development caused defects in NC-derived tissues, but we lacked an understanding of the mechanism through which NSAIDs alter NC development and the developmental window during which NSAID-specific effects occur. Our data shows that genes encoding the COX enzymes and other key effector molecules in the signaling pathway appear to be expressed at the right place and time to play a role in early NC development, including during NC specification and migration (Figures 1-3)(Leathers et al., 2024a). We found that global COX signlaing inhibition through exposure to the NSAID, NPX, has significant effects on the gross morphological development of axolotl embryos, which may be attributed to abnormal development of NC and CNS cells, among other tissues (Figure 9). We identified significant changes in the number and migratory capacity of both SOX9+ and PAX7+ NC cells in tailbud embryos, suggesting a role for COX signaling during early NC cell migration, specifically in SOX9+ cells fated to become chondrocytes in the developing craniofacial skeleton (Figures 5, 6, 9). In disrupting normal SOX9 signaling, we found that COL2A1-expressing cells are capable of populating the developing lower jaw, but cartilage structures with normal morphology at the cellular and tissue level do not form (Figure 6, 9). Interestingly, in the NC cell GRN, SOX9 is not only an important regulator of chondrogenic differentiation in cranial NC cells, but it is also important for initiating the expression of SOX10 and the developmental program for the glial and pigment cell lineages of NC cells (Betancur et al., 2010; Carney et al., 2006; Lefebvre et al.; Zhao et al., 1997). We hypothesize that these regulatory relationships explain our data showing that cartilage is not the only NC derivative that appears to be affected by NPX exposure, evidenced by abnormalities seen in ectodermal derivatives of the PNS and CNS (Figures 7, 8). The NC-derived sensory structures of the axolotl skin also develop abnormally after exposure to NPX and show abnormal expression of ECAD and PAX7 demonstrating potential changes in cell potential and fate as a result of COX inhibition (Figure 87). Significantly reduced expression of TUBB3 and GFAP indicate to us that the effects of NPX on development may extend beyond NC cells to include neurodevelopmental effects as well (Figure 8).

**Figure 9.**
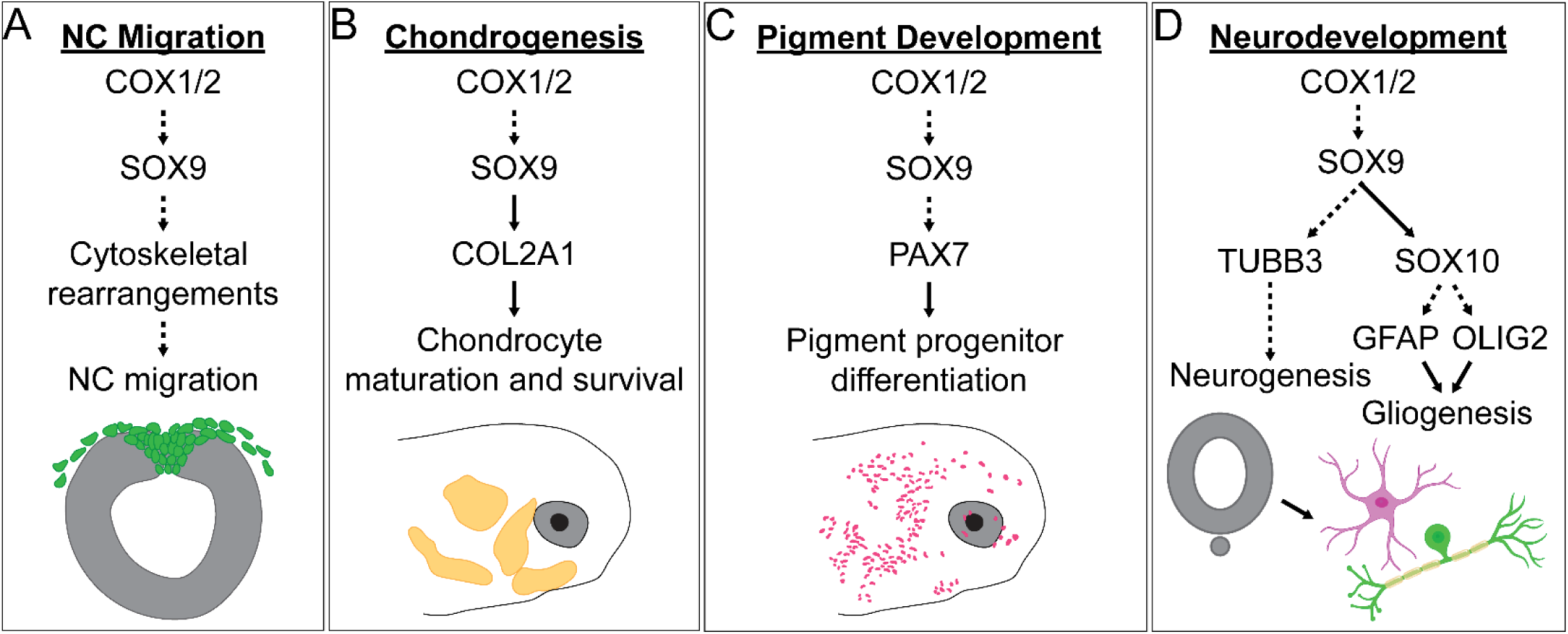
Graphical summary of the potential networks downstream of COX signaling driving development. Overview of the proposed mechanisms downstream of COX signaling with regard to (A) NC migration, (B) chondrogenesis, (C) pigment cell development, and (D) neurodevelopment based on prior work in other models. Solid black arrows indicate established signaling pathways or protein interactions. Dashed arrows highlight possible interactions that could serve as the basis for future studies. Astrocyte and neuron graphical icons were created using BioRender.

### COX signaling and SOX9-dependent cartilage and bone development

Our data suggest that COX signaling is necessary for the formation of bonafide NC cells and for NC cell migration based on altered PAX7 and SOX9 expression after NPX exposure (Figure 5). These results are in alignment with previous studies which demonstrated that exposure to NSAIDs inhibits NC cell migration, leading to abnormal development of NC-derived enteric nerves, but they suggest a homeostatic role for COX signaling during development in controlling SOX9 expression and cell migration rather than the classical role of the COX signaling pathway in adults as a mediator of inflammation. As SOX9 is expressed in many tissues throughout the developing embryo and is necessary for several key developmental processes including skeletogenesis, nephrogenesis, and gonadogenesis, it is reasonable to conclude that disruption in SOX9 signaling following NSAID exposure may also be the cause of other morphological defects that were not investigated here as they were beyond the scope of this study (Barrionuevo and Scherer, 2010; Cheung and Briscoe; Jo et al., 2014; Mori-Akiyama et al., 2003).

With respect to the potential role of COX signaling in NC cell migration, there is exisiting evidence to support the hypothesis that NPX-mediated COX inhibition alters directional NC cell migration through disruption of Rac and Rho activation and signaling. During the epithelial to mesenchymal transition (EMT) which preceedes early NC migration, NC cell polarization changes as they delaminate, collectively migrate, and become mesenchymal and invasive. Prior work from our lab and others identified that during NC EMT, there are rapid changes in cell adhesion proteins and actin polymerization (Matthews et al., 2008; Rogers et al., 2013). These changes establish an expression gradient of small G protiens like Rac1 and RhoA which are involved in essential downstream signaling pathways necessary for normal cell migration and tissue morphogenesis (Mayor and Theveneau, 2014; Sasaki et al., 2010; Suzuki et al., 2009; Tang et al., 2022; Yalovsky et al., 2008). Since prostaglandins can induce activation of Rac1 via siganling through EP3 (Hatae et al., 2002; Katoh et al., 1996), it is possible that these pathways may be linked and that inhibition of COX signaling via NSAID exposure may prevent the necessary cell adhesion and cytoskeletal arrangement changes downstream of SOX9 activation that facilitate establishment of Rac/Rho gradients within the cell causing reduced cell migration (Figure 9A).

In NPX-exposed embryos, COL2A1-expressing cells populate the developing lower jaw but are incapable of creating stereotypical cartilage structures with normal morphology at the cellular and tissue level (Figure 6). We propose that the change in SOX9 expression is sufficient to alter expression, potentially via a developmental delay, of COL2A1 in early development of the cranial cartilages (Lefebvre et al., 1997; Zhao et al., 1997). In addition to regulating COL2A1 expression, SOX9 has previously been linked to several steps in cartilage development including mesenchymal condensation, chondrocyte differentiation and maturation (Akiyama et al., 2002; Zheng et al., 2024). Prior studies identified that SOX9 is essential for maturation and survival of chondrocytes through its interactions with the PI3K/Akt pathway— a cononical regulatory pathway of cell growth, survival, and proliferation (Ikegami et al., 2011). Cross-talk between SOX9 signaling and the PI3K/Akt pathway has previously been studied in various contexts including cancer cell survival and proliferation, and tissue regeneration and fibrosis, which may provide useful insight into future research in the context of embryonic development (Figure 9B) (Chen et al., 2021; Zhang et al., 2020).

Our data recapitulates abnormal craniofacial phenotypes following NSAID exposure in early development with similarity to the mandibular hypoplasia phenotype often observed in disorders associated with mutations in the *SOX9* gene such as campomelic dysplasia and Pierre Robin sequence (Al-Qattan and Almohrij, 2022; Csukasi et al., 2019; Hossain et al., 2023). These links highlight the need for further investigation into the dysregulation of SOX9 signaling in NSAID-mediated developmental toxicity, because other characteristics of SOX9-related disorders include abnormal limb development, sex determination, and gonadogenesis (Barrionuevo and Scherer, 2010; Cheung and Briscoe; Jo et al., 2014; Mori-Akiyama et al., 2003). Additionally, Meckel’s cartilage typically has distinct domains across the anterior-posterior axis which uniquely contribute to development of the adult mandible, supporting structures like the mandibular symphysis, and bones of the middle ear, so disruption of its formation throughout the cranial region could has significant consequences for the establishment of these domains and ultimately their contributions to the developed mandibular structural components (Pitirri et al., 2022; Svandova et al., 2020).

### COX signaling and PAX7-mediated pigment development

SOX9 initiates the expression of SOX10 in the NC GRN, which drives the developmental program for the pigment cell, glial, and neuronal lineages of NC cells (Betancur et al., 2010; Carney et al., 2006; Lefebvre et al., 1997; Zhao et al., 1997). Based on our studies, the NC-derived sensory structures and pigment cells in the axolotl skin develop abnormally after exposure to NPX and exhibit abnormal expression of ECAD and PAX7 demonstrating possible changes in cell potential and fate as a result of COX inhibition (Figure 7). PAX7 expression is essential in regulating formation of the melanophore and xanthophore lineages of pigment cells (Miyadai et al., 2023a; Nord et al., 2016), so altered PAX7 expression downstream of the effects on SOX9 could result in abnormal development of these pigment cell types (Figure 9C). Specifically, abnormal pigment cell development could occur due to changes in expression of known PAX7 targets including *SOX10* in early NC migration, and later, *MITF* (Miyadai et al., 2023a; Miyadai et al., 2023b; Nord et al., 2016). However, although historical studies on xanthophore development in axolotls exist (Thibaudeau and Holder, 1998; Thibaudeau et al., 1998), there are many gaps in our understanding of the genetic and molecular regulation of pigment cell development beyond chromatophores in axolotl morphs outside of the melanoid color variant (Kabangu et al., 2023). Future research should focus on identifying the key regulators of this process and the exact role of PAX7 in the decision between melanophore and xanthophore formation from NC progenitors.

Beyond its role in pigment cell development, PAX7 is also a key regulator of myogenesis during embryonic development (Buckingham and Relaix, 2015; Musumeci et al., 2015; Otto et al., 2006). The dysregulation of PAX7 expression following NSAID-mediated COX inhibition could therefore further explain the changes in axial development that we observed (Figure 4).

### COX signaling and PNS and CNS development

There is much to be discovered about the mechanisms driving development of lateral line sensory organs in the axolotl, but based on previous studies in zebrafish, we know their development requires precise patterns of cell migration, proliferation, and specific cues to differentiate in order to develop normally (Laguerre et al., 2005; Thomas et al., 2015; Wada et al., 2013). Examples of necessary cues include interactions with developing glial cells in peripheral tissues and normal innervation by placodally- and NC-derived sensory neurons (Grant et al., 2005; Northcutt and Brandle, 1995; Schlosser and Northcutt, 2001; Schuster et al., 2010; Wada et al., 2013). A potential reduction of SOX10 expression due to reduced SOX9 expression following NSAID exposure would likely have effects on the development of glial and neuronal cells in the periphery, which we observed as reduced peripheral TUBB3 and GFAP expression in embryos exposed to NPX (Figure 8). This putative perturbation of SOX10 expression may impair interactions between developing glial cells and the migrating sensory cell precursors, resulting in abnormal lateral line sensory organ development (Figure 9D).

We also identified concentration-dependent abnormalities in expression of markers of neurons and glia in the ectodermally-derived CNS. Significantly reduced expression of TUBB3 and GFAP indicate to us that the effects of NPX exposure on development may extend beyond NC cells to include the developing CNS as well (Figure 8). Previous studies have proposed a role for SOX9 in the processes of glial cell fate determination in neuroepithelial progenitor populations and cell proliferation during neocortical expansion which may explain this observed phenotype, but further analyses would be required to definitively link NPX exposure to this phenoytpe in axolotls (Guven et al., 2020; Vogel and Wegner, 2021).

### Broader implications of study

Despite the established link between NSAID use and developmental abnormalities, it is reported that one in four women take NSAIDs during pregnancy with 5% of these women using over-the-counter NPX for pain relief (Werler et al., 2005). Additionally, NSAID use is not explicitly contraindicated in pregnant women until after week twenty of pregnancy, once significant NC development is already complete (O’Rahilly and Muller, 2007). Although pain and inflammation are common complications of pregnancy, we still have a poor understanding of the potential risks of NSAID treatment during prenancy, and as such, it is imperative to understand the effects of NSAID exposure so that safe and effective treatment plans using these therapeutics can be devised for use in medicine. Here, we have identified a link between inhibition of COX signaling through NSAID exposure and abnormal NC cell development which will serve as the premise for future mechanistic research into the developmental necessity of COX signaling and potential teratogenicity of NSAIDs, ultimately informing decision-making regarding pain management during pregnancy. Overall, further research into the developmental toxicity of NSAIDs will allow clinicians to perform more accurate risk assessment when recommending NSAID use to pregnant patients.

## V. AUTHOR CONTRIBUTIONS

Conceptualization: CDR and EJM. Data Curation (experiments and imaging): EJM, TL, NMW, KS, MK, and RR. Formal Analysis: EJM and RR. Funding Acquisition: CDR, EJM, and RR. Methodology: CDR, EJM, and RR. Writing: CDR, EJM, and RR. Supervision: CDR.

## VI. ACKNOWLEDGEMENTS

The authors would like to acknowledge the following funding sources: NSF CAREER award 2143217 and NIH R03DE032047-01 to CDR. Funding for EJM was provided by the UC Davis eMCDB T32 (GM-153586) and Research Corporation for Science Advancement (Scialog ABI), and funding for RR was provided by the UC Davis Vision Sciene T32 (NEI-EY015387). We would like to thank the members of the Rogers Lab at UC Davis and the UC Davis community for their discussions and input on this project.

## VII. Supplemental Materials

**Supplemental Figure 1.**
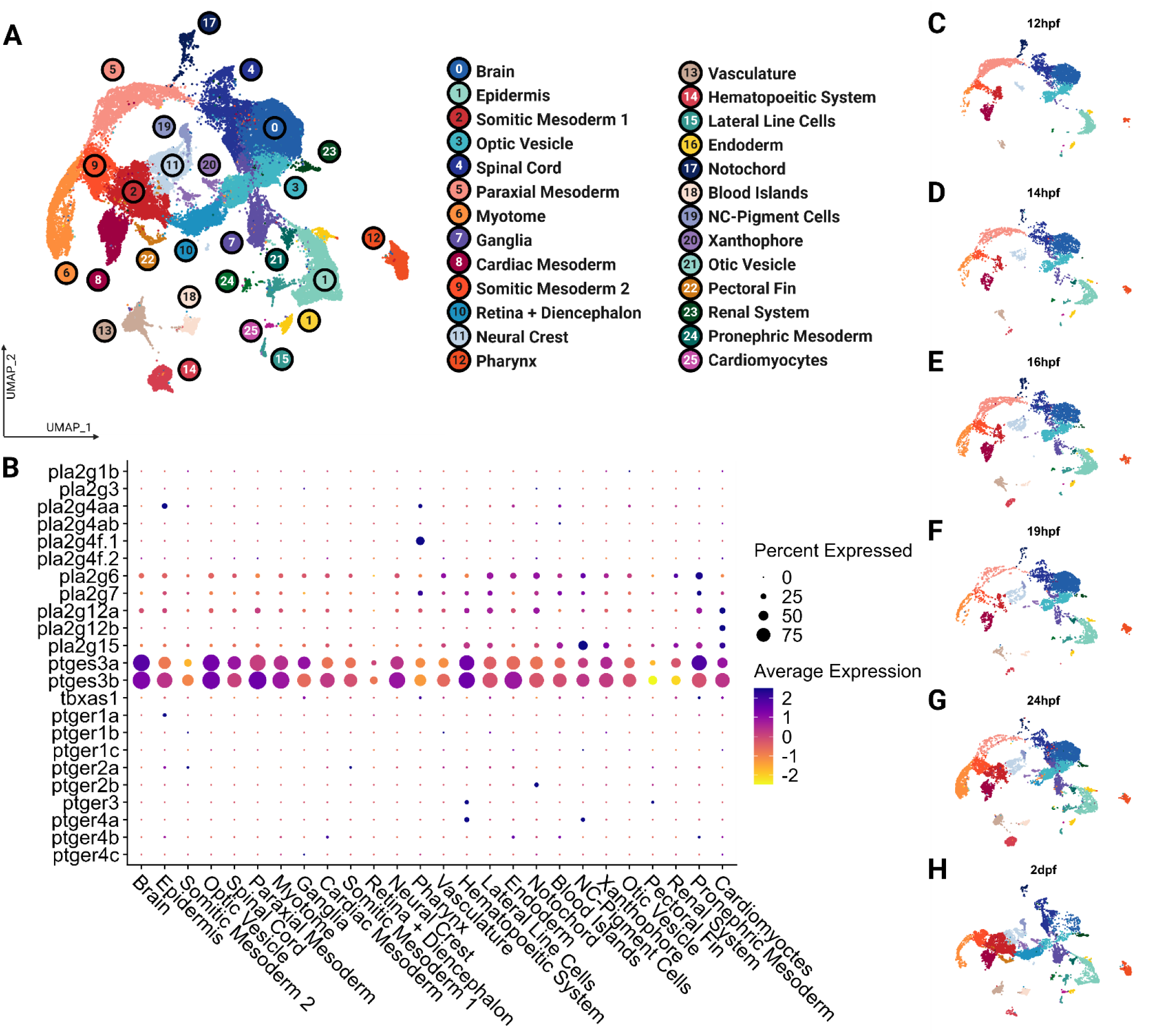
Expression of cyclooxygenase enzymes across zebrafish cell types. Unsupervised clustering of publicly available scRNA-seq data of zebrafish embryos between 12hpf-2dpf demonstrates expression of cyclooxygenase enzymes. (A) UMAP demonstrating the unsupervised clustering results of zebrafish embryos. (B) Dot plot demonstrating expression levels of cyclooxygenase pathway members across zebrafish cell types. (C-H) UMAP split by stages. NC, neural crest; hpf, hours post-fertilization; dpf, days post-fertilization.

**Supplemental Figure 2.**
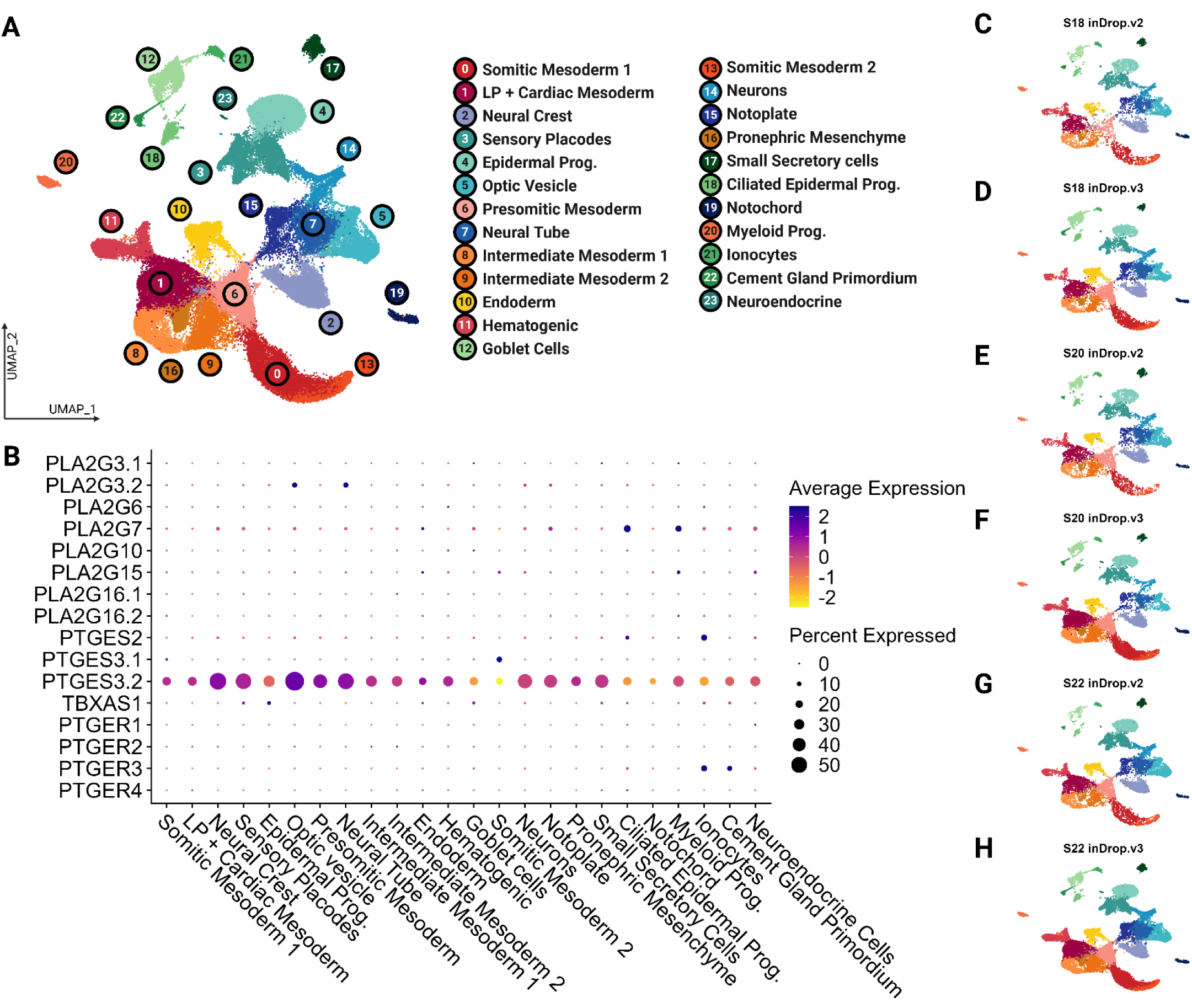
Unsupervised clustering of publicly available scRNA-seq data of African clawed frog embryos between stage 18 and 22 demonstrates expression of cyclooxygenase enzymes. (A) UMAP demonstrating the unsupervised clustering results of frog embryos. (B) Dot plot demonstrating expression levels of cyclooxygenase pathway members across frog cell types. (C-H) UMAP split by stage and inDrop version. LP, lateral plate; Prog., progenitors.

**Supplemental Figure 3.**
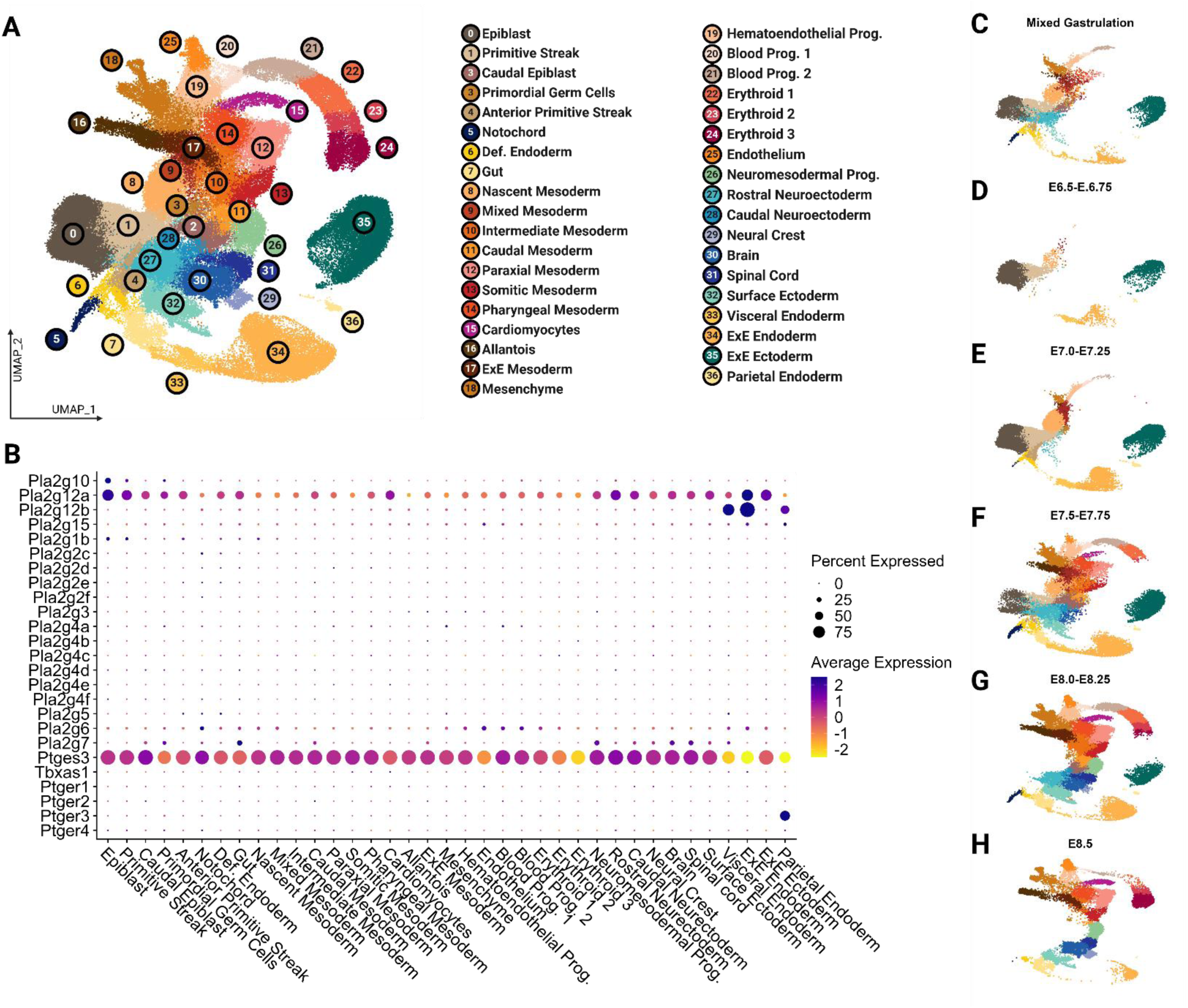
Expression of cyclooxygenase enzymes across mouse cell types. Unsupervised clustering of publicly available scRNA-seq data of house mouse embryos between late gastrulation and E8.5 demonstrates expression of cyclooxygenase enzymes. (A) UMAP demonstrating the unsupervised clustering results of mouse embryos. (B) Dot plot demonstrating expression levels of cyclooxygenase pathway members across frog cell types. (C-H) UMAP split by stage. Def, definitive; Prog., progenitors; ExE, extraembryonic.

**Supplemental Figure 4.**
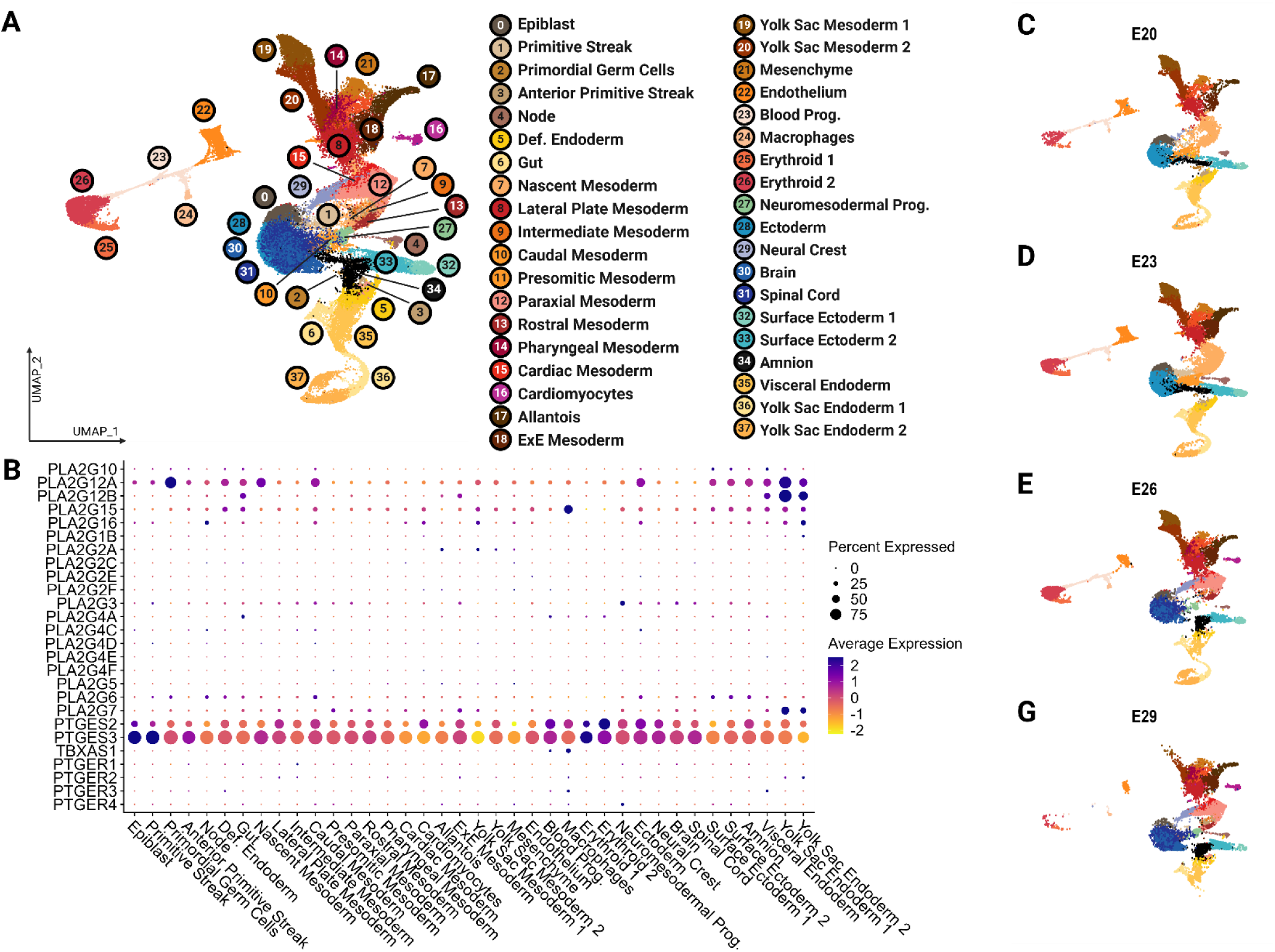
Expression of cyclooxygenase enzymes across macaque cell types. Unsupervised clustering of publicly available scRNA-seq data of Cynomolgus macaque embryos between E20 and E29 demonstrates expression of cyclooxygenase enzymes. (A) UMAP demonstrating the unsupervised clustering results of macaque embryos. (B) Dot plot demonstrating expression levels of cyclooxygenase pathway members across macaque cell types. (C-H) UMAP split by stage. Def, definitive; Prog., progenitors; ExE, extraembryonic.

**Supplemental Figure 5.**
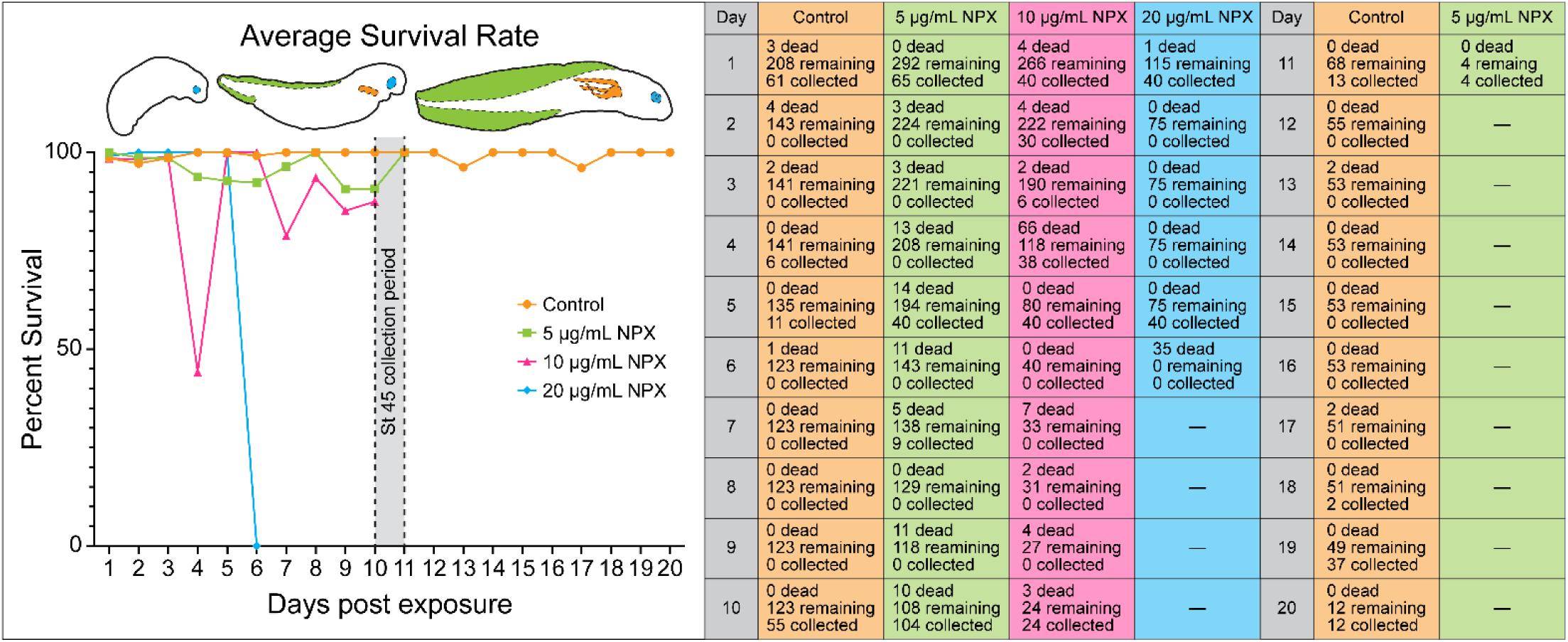
NSAID exposure in early development reduces axolotl embryo survival in a dose-dependent manner. (Left) Embryos treated with NPX show reduced survival rate compared to control embryos. To calculate the survival rate per day the difference between the number of embryos remaining on day n and those that died between day (n-1) and day n was first calculated (# remaining - # dead). This number was then divided by the number of embryos remaining on day n and multiplied by 100 to get a percentage that survived between the two days of interest [(# remaining - # dead)/# remaining x100]. Dashed outline highlights the collection period for stage 45 embryos. Embryo drawings shown to demonstrate the normal morphological development of axolotl embryos. (Right) Numbers of embryos that were either found dead, were alive and apparently healthy, or were collected for bright-field imaging and IHC experiments based on the number of days post exposure to NPX.

**Supplemental Figure 6.**
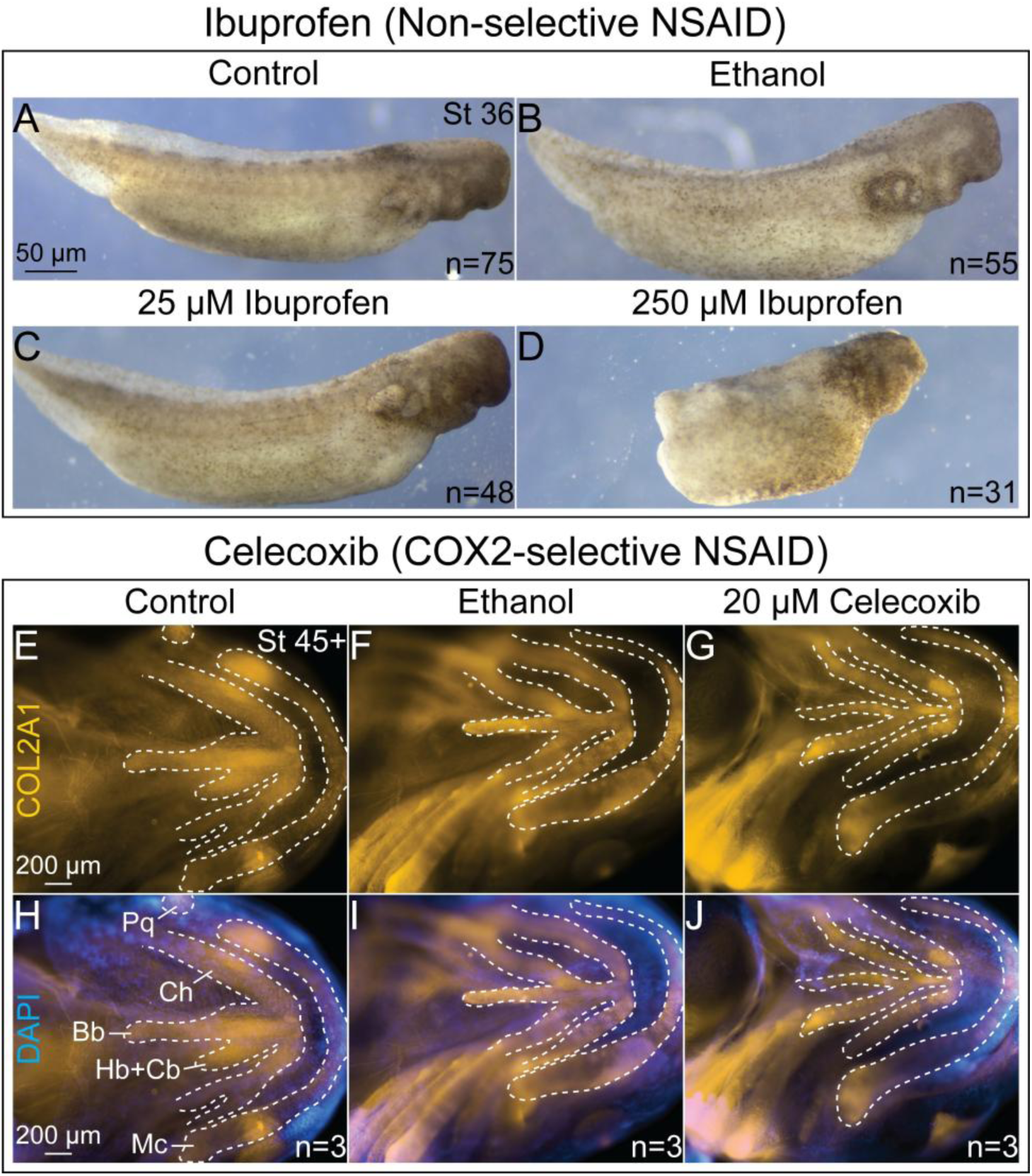
COX-2-selective NSAIDs do not cause the craniofacial defects observed in embryos exposed to non-selective NSAIDs. (A-D) Bright-filed images of stage 36 axolotl embryos demonstrating that high dose exposure (250 µM) to the non-selective NSAID, Ibuprofen, causes significant axial and craniofacial defects similar to those seen in NPX treated embryos. (E-J) Wholemount IHC images for COL2A1 taken ventrally to highlight the apparently normal development of cranial cartilage structures in stage 45 tadpoles exposed to the COX-2-selective NSAID, Celecoxib. Bb= basibranchial cartilage, Cb= ceratobranchial cartilage, Ch= ceratohyal cartilage, Hb= hypobranchial cartilage, Mc= Meckel’s cartilage, Pq= palatoquadrate cartilage.

**Supplemental Figure 7.**
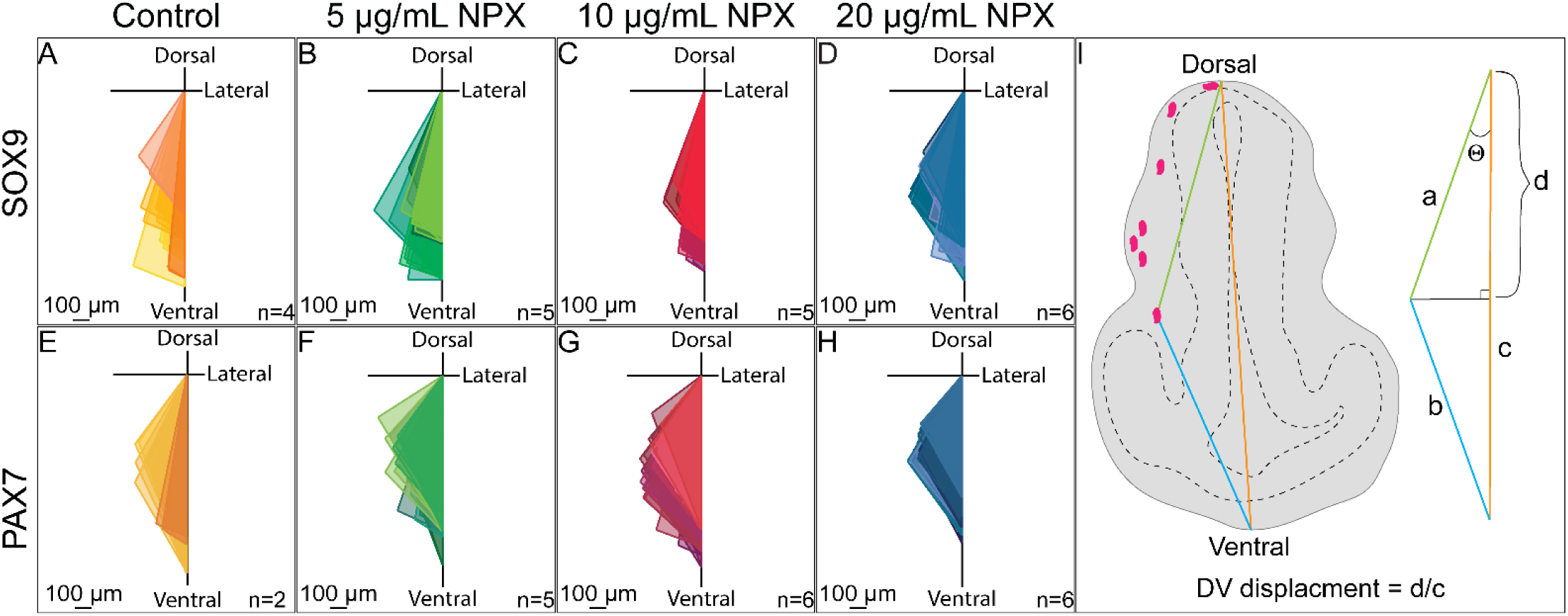
Quantification of the DV displacement of SOX9/PAX7+ NC cells in the cranial region. Triangles drawn in Adobe Illustrator showing the migration patterns of (A-D) SOX9+ or (E-H) PAX7+ NC cells in transverse sections of stage 28 axolotl embryos. All triangles of the same color are from sections of the same embryo. (I) Graphical representation of a transverse section of a stage 28 axolotl embryos demonstrating the measurments and calculations used to determine the DV displacement of NC cells for each section image (quantified in Figure 4R).

**Supplemental Table 1.** Methods for single cell analysis of publicly available embryonic data.

| Figure | Species | Data Set Features | Data Set Processing | Source |
| --- | --- | --- | --- | --- |
| Figure 1A-G, Supp. Figure 1 | *Danio rerio* (Zebrafish) | Whole embryo (12hpf, 14hpf, 16hpf, 19hpf, 24hpf, 2dpf) | Stage specific had files converted to *Seurat* v5 objects. Objects were integrated using *harmony* as per *Seurat* integration pipeline (Hao et al., 2024; Korsunsky et al., 2019). Cluster annotation was determined using existing annotations from source manuscript and top marker genes of each cluster. | Datasets obtained from Zebrahub (Farnsworth et al., 2020; Farrell et al., 2018; Raj et al., 2020; Wagner et al., 2018). https://zebrahub.sf.czbiohub.org/transcriptomics |
| Figure 1H-N, Supp. Figure 2 | *Xenopus laevis* (African clawed frog) | Whole embryo (Stage 18, 20, 22) | The count matrix and metadata files were converted to *Seurat* objects. Genes were renamed for analysis and can be made available on request. Objects were integrated using *harmony* as per *Seurat* integration pipeline [132, 133]. Utilized 30 dimensions for reduction and 0.5 resolution for k-means clustering. Cluster annotation was determined using existing annotations from source manuscript and top marker genes of each cluster. | Datasets obtained from Tabula Rana: *Xenopus* Embryo Single Cell Atlas (Petrova et al., 2024) https://xenopus.hms.harvard.edu/Embryo.html |
| {Figure 1O-U, Supp. Figure 3 | *Mus musculus* (House mouse) | Whole embryo (E6.5, 6.75, E7.0, E7.25, E7.5, | The processed *SingleCellExperiment* object was downloaded via R package *MouseGastrulationData* (https://www.bioconductor.org/packages/release/data/experiment/html/MouseGastrulationData.html). Object was converted to a *Seurat* v5 object, | (Pijuan-Sala et al., 2019) |
|  |  | E7.75,<br>E8.0,<br>E8.25,<br>E8.5) | normalized, and scaled. Dimensional reduction coordinates and cell type annotations were maintained from source manuscript. Cell type colors were reassigned. |  |
| Figure 1V-AB, Supp. Figure 4 | *Macaca fascicularis*<br>(Cynomolgus monkey) | Whole embryo (CS8, CS9, CS11) | The filtered feature matrix was obtained from NCBI GEO (GSE193007) and converted to a *Seurat* v5 object, normalized, and scaled. Metadata, including cell type annotation, and dimensional reduction coordinates were maintained from source manuscript. Cell type colors were reassigned. | (Zhai et al., 2022) |

